# Single Cell Resolution Tracking of Cutaneous T-Cell Lymphoma Reveals Clonal Evolution in Disease Progression

**DOI:** 10.1101/2025.02.11.637715

**Authors:** Hannah K. Dorando, Jared M. Andrews, Nicholas C. Borcherding, Chaz C. Quinn, Jennifer A. Schmidt, Oam U. Khatavkar, Jahnavi Aluri, Michael T. Harmon, Marcus P. Watkins, Anastasia Frank, Megan A. Cooper, Amy C. Musiek, Neha Mehta-Shah, Jacqueline E. Payton

## Abstract

Cutaneous T-cell lymphoma (CTCL) remains a challenging disease due to its significant heterogeneity, therapy resistance, and relentless progression. Multi-omics technologies offer the potential to provide uniquely precise views of disease progression and response to therapy. We present here a comprehensive multi-omics view of CTCL clonal evolution, incorporating exome, whole genome, epigenome, bulk-, single cell (sc) VDJ-, and scRNA-sequencing of 114 clinically annotated serial skin, peripheral blood, and lymph node samples from 35 CTCL patients. We leveraged this extensive dataset to define the molecular underpinnings of CTCL progression in individual patients at single cell resolution with the goal of identifying clinically useful biomarkers and therapeutic targets. Our studies identified a large number of recurrent progression-associated clonal genomic alterations; we highlight mutation of CCR4, PI3K signaling, and PD-1 checkpoint pathways as evasion tactics deployed by malignant T cells. We also identified a gain of function mutation in STAT3 (D661Y) and demonstrated by CUT&RUN-seq that it enhances binding to transcription start sites of genes in Rho GTPase pathways, which we previously reported to have activated chromatin and increased expression in HDACi-resistant CTCL. These data provide further support for a previously unrecognized role for Rho GTPase pathway dysregulation in CTCL progression. A striking number of progression-associated mutations occurred in chromatin methylation modifiers, including EZH2, suggesting that EZH1/2 inhibition may also benefit patients with CTCL. Knowledge of these molecular changes should be leveraged for improved disease monitoring, biomarker-informed clinical trial design, and new therapeutic strategies in this challenging and incurable cancer.

## INTRODUCTION

Cutaneous T-cell Lymphomas (CTCL) are a heterogeneous subset of T-cell Non-Hodgkin Lymphoma that often involve skin, lymph nodes, and blood. Though early-stage patients have an excellent prognosis, patients with advanced-stage CTCL (stages ≥IIB) have a 5-year survival of only 20%^1^. Few therapies lead to long-lasting remissions and therapeutic decision-making guidance is lacking. Furthermore, little is known regarding disease evolution on a genomic level. Therefore, we reasoned that defining biomarkers of CTCL progression will identify new therapeutic targets and provide guidance for therapy selection at times of progression.

Previous studies, including our own, have reported recurrent molecular changes in the genomes, epigenomes, and transcriptomes of CTCL^2–14^. Notably, most of these examined patients at a single point in time, leaving the molecular drivers of disease progression and therapy resistance undefined. In contrast, our previous study^2^ evaluated chromatin histone acetylation and gene expression in serial samples from CTCL patients before, during, and after initiation of histone deacetylase inhibitor therapy (HDACi). We identified enhancer deregulation and associated transcriptional changes in CTCL tumors that were resistant to HDACi. Deregulation was enriched in pathways involved in cell adhesion and migration, cell cycle/mitosis, Rho GTPase signaling, and inflammatory response^2^. These compelling findings led us to ask whether deregulation of these pathways is specific to HDACi resistance or rather is more broadly involved in therapy resistance and/or disease progression.

To address this knowledge gap, we applied complementary multi-omics approaches to serial samples collected from the same CTCL patients over time. Multi-omics and single cell technologies can provide insights regarding disease progression and response to therapy. We present here the results of our study that is, to our knowledge, the most comprehensive multi-omics study to date, incorporating exome, whole genome, epigenome, and bulk RNA-, single cell (sc)VDJ- and scRNA-sequencing of 114 samples from 35 CTCL patients. We leveraged this unique, clinically annotated multi-omics dataset to define the clonal evolution and temporal progression of CTCL at single cell resolution. Our results identified somatic genome copy number alterations (CNA) and single nucleotide variants (sSNV) that are recurrent across patients. While CNA were more frequent, consistent with other reports^6,10^, we found many genes harboring both CNA and sSNV, suggesting that these play key roles in CTCL pathogenesis. More compelling is our finding that some of these alterations are acquired with progression or therapy resistance (APR), including a striking number of chromatin methylation modifiers. Our results present a comprehensive multi-omics view of clonal evolution in CTCL and reveal molecular changes that are acquired with disease progression and therapy resistance, presenting new opportunities for therapeutic intervention in this challenging cancer.

## METHODS

### Sample collection

De-identified peripheral blood draws, skin punch, or lymph node biopsies were obtained from patients seen at the Washington University School of Medicine Cutaneous Lymphoma Clinic under IRB-approved protocols with patients providing informed consent between 2015-2019.

### Exome + whole genome, bulk and single cell RNA sequencing

Exome + whole-genome sequencing (eWGS) was performed on DNA isolated from purified malignant T cells or matched non-malignant cells from peripheral blood, or from frozen or FFPE biopsies (skin or lymph node); in total, there were 26 samples from ten CTCL patients. Bulk RNA sequencing was performed on 68 samples from 35 CTCL patients and 5 healthy controls. Single cell RNA and TCR VDJ RNA sequencing was performed on 16 samples from 6 CTCL patients.

*Additional methods are provided in Supplemental Methods*.

## RESULTS

### Study overview, patient demographics and sample collection

To map the evolution of CTCL genomes and transcriptomes over time, we collected 114 peripheral blood (PB), skin, and lymph node (LN) biopsy specimens from 35 CTCL patients seen at the Washington University School of Medicine Cutaneous Lymphoma Clinic and 5 healthy donors (**Figure 1A**, **Table 1**). For the multi-omics cohort, half of first specimens were collected within 60 days of diagnosis with subsequent specimens collected at progression; there was an average of 20 months between collections (1.4 – 60, SD = 11.8) with 4-11 serial samples per patient (**Figure 1B**). Patients received a median of one therapeutic regimen (range 1 - 9) during the period of specimen collection.

**Figure 1.**
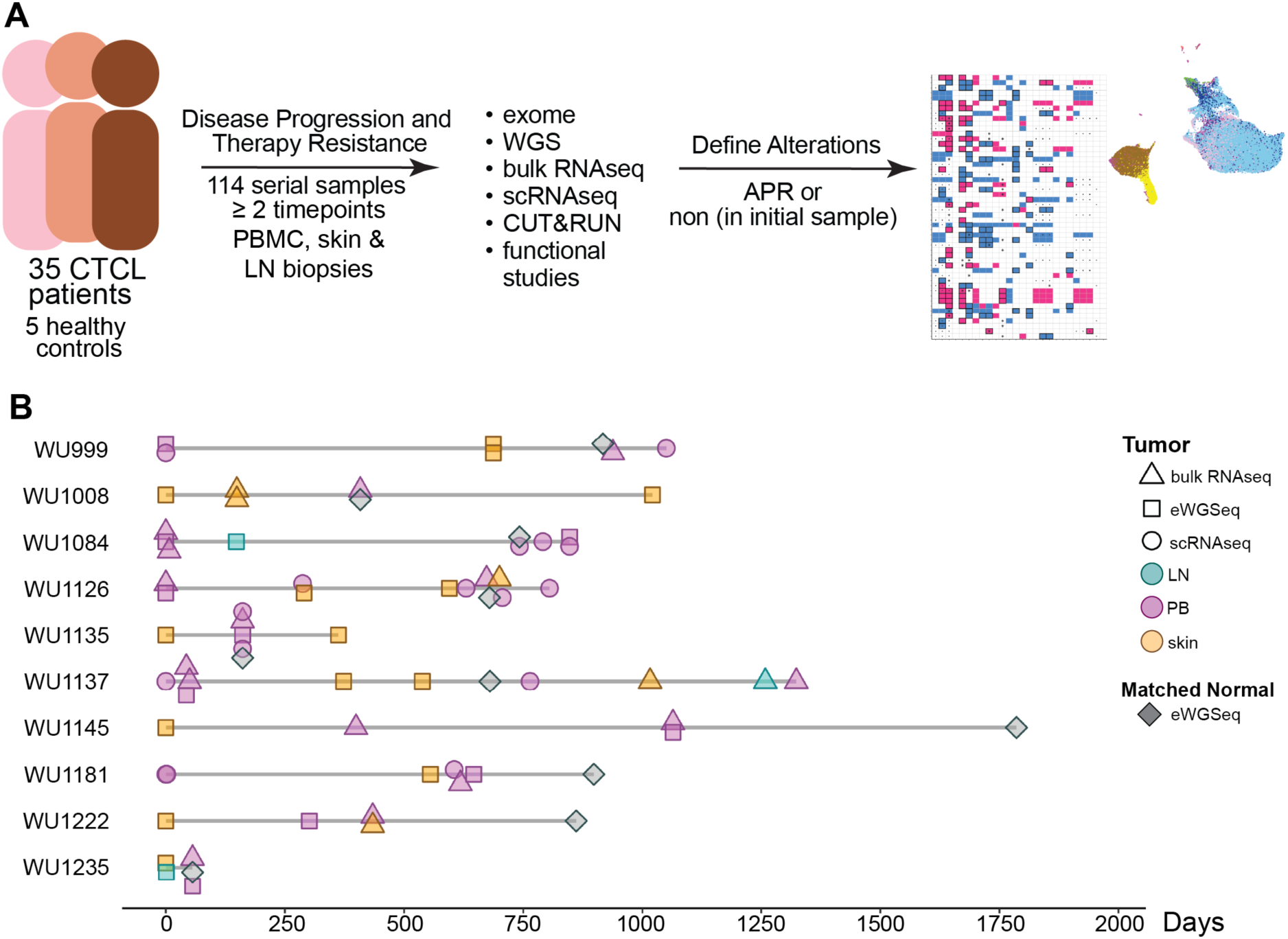
Study overview. **A)** Diagram depicting the overall study. A total of 114 peripheral blood (PB), skin and lymph node (LN) biopsy specimens were collected from 35 MF/SS patients and 5 healthy donors. For 10 of the MF/SS patients, serial specimens (4-11 per patient), including matched non-malignant cells, were collected. Bulk RNAseq was performed on at least 1 sample from all individuals. Specimens from the cohort of 10 patients were additionally subjected to single cell (sc) RNA-seq, and exome + whole genome sequencing (eWGS) to define expression profiles, map cell clusters and TCR clonality, and identify single nucleotide and copy number variants present in tumor but not matched non-malignant cells. The functional impact of a somatic single nucleotide variant identified in STAT3 was evaluated using CUT&RUN-seq and gene regulation assays. **B)** Timeline graph of serial specimens collected for each patient. Colors indicate specimen type and shapes indicate ‘omics assay type.

**Table 1.** Patient Demographics.

|  | **Total cohort** | **Multi-omics cohort** |
| --- | --- | --- |
| **Sample size, n** |  |  |
| **Median age (range)** | 68 (39-88) | 64.5 (39-74) |
| **Female, n (%)** | 16 (46%) | 5 (50%) |
| **Race** |  |  |
| Caucasian/White, n (%) | 27 (77%) | 6 (60%) |
| AA or Black, n (%) | 8 (23%) | 4 (40%) |
| Asian, n (%) | 0 | 0 (0%) |
| Unknown, n (%) | 0 | 0 (0%) |
| **Diagnosis** |  |  |
| CTCL, n (%) | 1 (3%) | 0 (0%) |
| Cutaneous gamma-delta T-cell lymphoma, n (%) | 1 (3%) | 0 (0%) |
| MF, n (%) | 12 (34%) | 3 (30%) |
| MF - transformed during treatment, n (%) | 4 (11%) | 0 (0%) |
| MF - transformed at diagnosis, n (%) | 2 (6%) | 0 (0%) |
| Sezary, n (%) | 13 (37%) | 7 (70%) |
| Subcutaneous Panniculitis-like T-cell Lymphoma, n (%) | 1 (3%) | 0 (0%) |
| T-cell PLL, n (%) | 1 (3%) | 0 (0%) |
| **Treatment history** |  |  |
| Prior # of therapies, median (range) | 1 (0-9) | 1.5 (0-6) |
| # of therapies during sample collection period, median (range) | 1 (1-9) | 3 (1-9) |
| Mogamulizumab | 12 (34%) | 7 (70%) |
| Mogamulizumab response | 8 (67%) | 4 (57%) |
| Check point inhibitors | 10 (29%) | 5 (50%) |
| Checkpoint inhibitor response | 6 (6%) | 3 (60%) |
| PI3K inhibitors | 6 (17%) | 4 (40%) |
| PI3K inhibitor response | 1 (17%) | 0 (0%) |
| HDACi | 22 (63%) | 8 (80%) |
| HDACi response | 8 (36%) | 3 (38%) |

### Serial WGS demonstrates both early and acquired genome copy number amplifications and deletions

We purified malignant T cells and non-malignant cells from a total of 36 samples from the multi-omics cohort; we confirmed purity by flow cytometry (**Figure S1)**. DNA extraction, whole genome sequencing and identification of copy number alterations (CNA) was performed as in ^15–17^ and **Figure S2**. We detected multiple large CNAs, up to entire chromosomes in size, in nearly all CTCL samples (**Figure 2A**), consistent with previous reports^6,10,14^.

**Figure 2.**
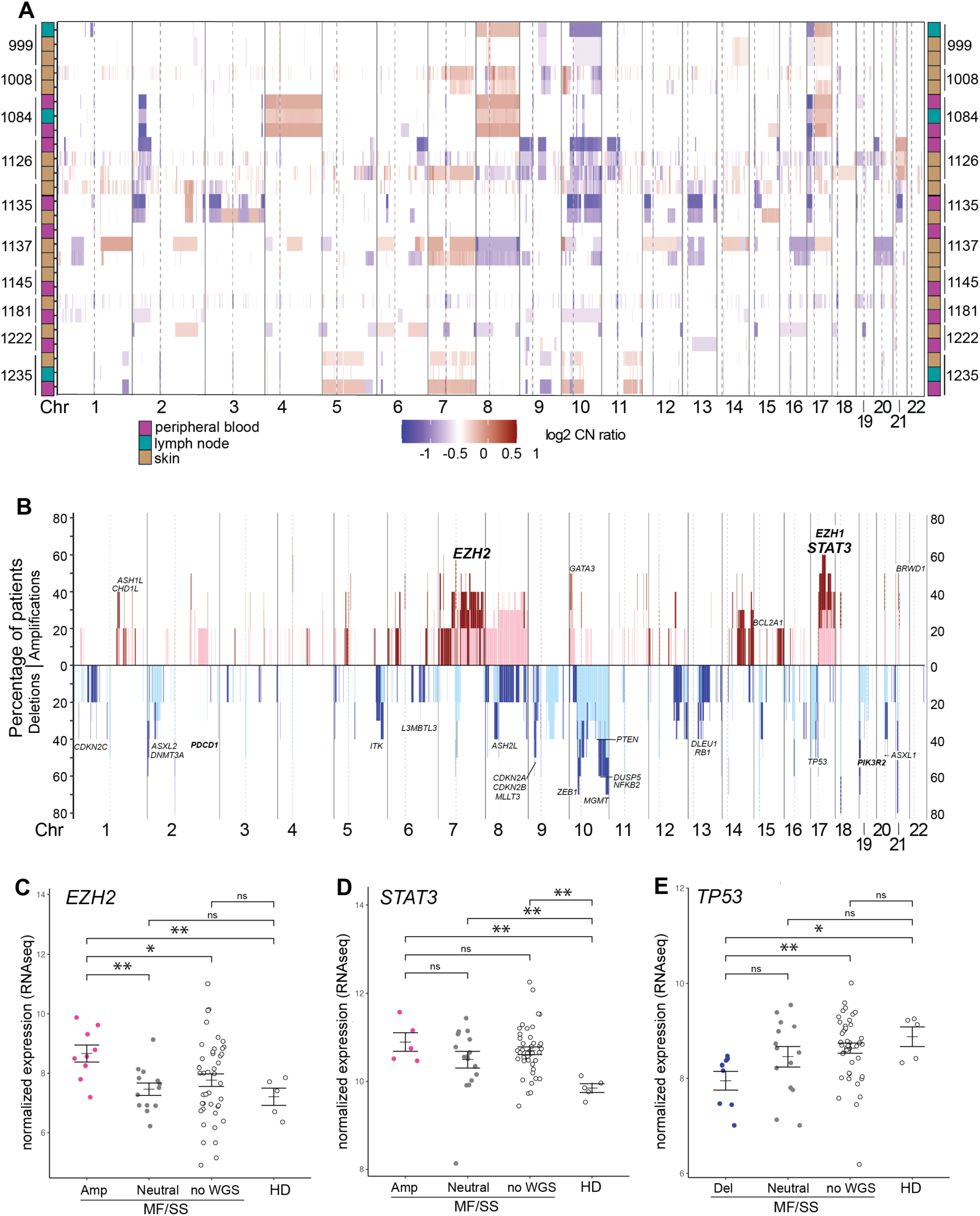
New copy number alterations are acquired with disease progression and correlate with gene expression. **A)** Whole genome log2 copy number (CN) ratio heat map shows data from each patient sample (rows) ordered by genomic location and chromosome. Copy number alterations (CNA) are colored according to their log2 copy number (CN) ratio of tumor:normal on a continuous color spectrum from blue (−1 to −0.2, deletion) to white (−0.2 to 0.2, no CNA) to red (0.2 to 1, amplification) in 100kb bins detected by WGS. Tissue types are color-coded for skin, peripheral blood, and lymph node indicated by color blocks to the left and right of the heatmap and are arranged in order of collection for each patient. **B)** Whole genome plot ordered by genomic location and chromosome shows the percentage of MF/SS patients with copy number amplification (above the horizontal line) or deletion (below the horizontal line) in 100kb bins detected by WGS. Lavender bars indicate copy number changes that were acquired with disease progression; gray bars indicate copy number changes that were present in the initial patient sample. **C-E)** Grouped dot plots depict normalized expression (bulk RNAseq) of example genes within CNA; CNA groups are based on WGS performed at the closest point in time to the RNAseq. HD – healthy donor. Wilcoxon rank sum test * p<0.05, ** p<0.01.

To identify CNAs that were gained or lost with disease progression, we compared CNAs in later samples to initial samples in matched patients. This analysis revealed a subset of CNAs that are <u>A</u>cquired with <u>P</u>rogression or <u>R</u>esistance to therapy (APR). **Figure 2B** shows the percentage of patients with APR gain or loss of each genomic region in lavender; CNAs present in the initial sample are in grey. Strikingly, many genes involved in epigenetic regulatory pathways are located within recurrent APR CNAs, including DNA and histone methyltransferases *DNMT3A*, *ASXL1/2*, *EZH1*, *EZH2*, and *KMT2C*, as well as chromatin remodelers *BRWD1*, *SMARCD2, SMARCE1*. Genes involved in cell cycle control such as *CDKN2A*, *CDKN2B*, and *CDKN2C* are lost with progression. *TP53* loss is present in the initial sample for most of the patients with this loss (4 of 5). Finally, APR CNAs contain several genes that code for important T-cell signaling molecules and transcription factors, including *GATA3*, *ITK*, *NFKB2*, *PTEN*, *STAT3*, and *ZEB1* (**Figure 2B**).

We next asked whether genes in CNAs were associated with corresponding changes in gene expression using bulk RNAseq from 35 CTCL patients and 5 healthy donors (68 total samples). We compared expression in CTCL samples with CNAs to CTCL samples with neutral copy number (determined by WGS from the closest timepoint) and to CTCL samples and healthy donors without known copy number (no WGS data available) (**Figures 2C-E, S3**). For some genes, expression significantly correlated with the “direction” of CNAs (i.e., CNA deletion – decreased expression, CNA amplification – increased expression). For example, samples with genomic amplification of *EZH2* exhibited significantly higher *EZH2* expression compared to those with neutral genome copy number and healthy donors. Amplified samples show a non-significant trend toward higher *EZH2* expression compared to samples with unknown copy number, which may be due to genome amplification in some of the samples without WGS data (**Figure 2C**). Genomic amplification of *EZH1* also demonstrated a non-significant trend toward higher expression levels (**Figure S3D**). Other genes also exhibited correlative changes in expression and genome copy number alterations: *STAT5A* and *BRWD1* (amplified) and *ITK, L3MBTL3*, *MGMT* and *PTEN* (deleted) (**Figure S3**). Notably, *STAT3* expression was significantly higher across CTCL samples compared to healthy donors and was not different between amplified and neutral samples, suggesting that multiple regulatory mechanisms cause high levels of *STAT3* expression (**Figures 2D, S3F**). Finally, *TP53* expression was the lowest with genomic deletion (**Figure 2E**). Together, these data reveal genes that are recurrently gained or lost in CTCL progression and whose correlative gene expression indicates a functional impact of genomic changes.

### APR somatic single nucleotide variants (sSNV) and CNA are enriched in epigenetic, DNA repair, and immune signaling pathways

The same malignant T cells and patient-matched non-malignant cells were subjected to whole exome sequencing and analysis as in ^15–17^ and **Figure S2**. Somatic (s)SNVs were subjected to read depth, artifact, annotation, and variant effect prediction filters. APR sSNVs were determined in the same way as APR CNAs. We identified 7353 genes that harbored at least one sSNV and at least one CNA across at least two patients. We used gene ontology and pathway tools to consolidate the genes into groups with common functions, then selected groups with the highest level of recurrent APR mutations for presentation in **Figure 3A**. The heatmap shows copy number, sSNV, and APR status of 50 of the genes with the most recurrent mutations; the mean of 51% demonstrates a high level of recurrence. Several of these genes demonstrate a significant correlation of CNA and expression (**Figures 2, S3**). Mutations in genes that code for proteins with epigenetic modifier function, especially DNA and histone methylation modifiers, were strikingly abundant. These included *ASXL1*, *EZH2*, *KMT2C*, *L3MBTL3,* and *NSD1,* all involved in epigenome methylation pathways (**Figures 3A-D, S3-S4**). As expected, genes in immune signaling pathways were recurrently mutated, including deletions and sSNV in the genes that code for IL2-inducible T-cell kinase (*ITK*), checkpoint factor PD-1 (*PDCD1*), phospholipase G1 (*PLG1*), and in the SH2 (p110 binding) and SH3 (PTEN binding) protein-interaction domains of PIK3R1 and PIK3R2, the regulatory subunits of phosphatidylinositol 3-kinase (PI3K) (**Figures 3A, E-F, S4A**). We identified CNA of *STAT3* and *STAT5A/B* in 70% of patients, with additional sSNVs in two patients. One patient harbored an APR sSNV in the SH2 domain of STAT3, D661Y, which has been reported as a somatic mutation in T-cell large granular lymphoma (T-LGL) and as a germline mutation in STAT3 GOF autoimmune syndrome^18,19^. We performed functional studies to further evaluate this mutation, see next section. Together, these analyses demonstrated that several key pathways, most prominently epigenome methylation, gain both CNA and sSNV with CTCL progression, supporting a role for these pathways in driving disease advancement.

**Figure 3.**
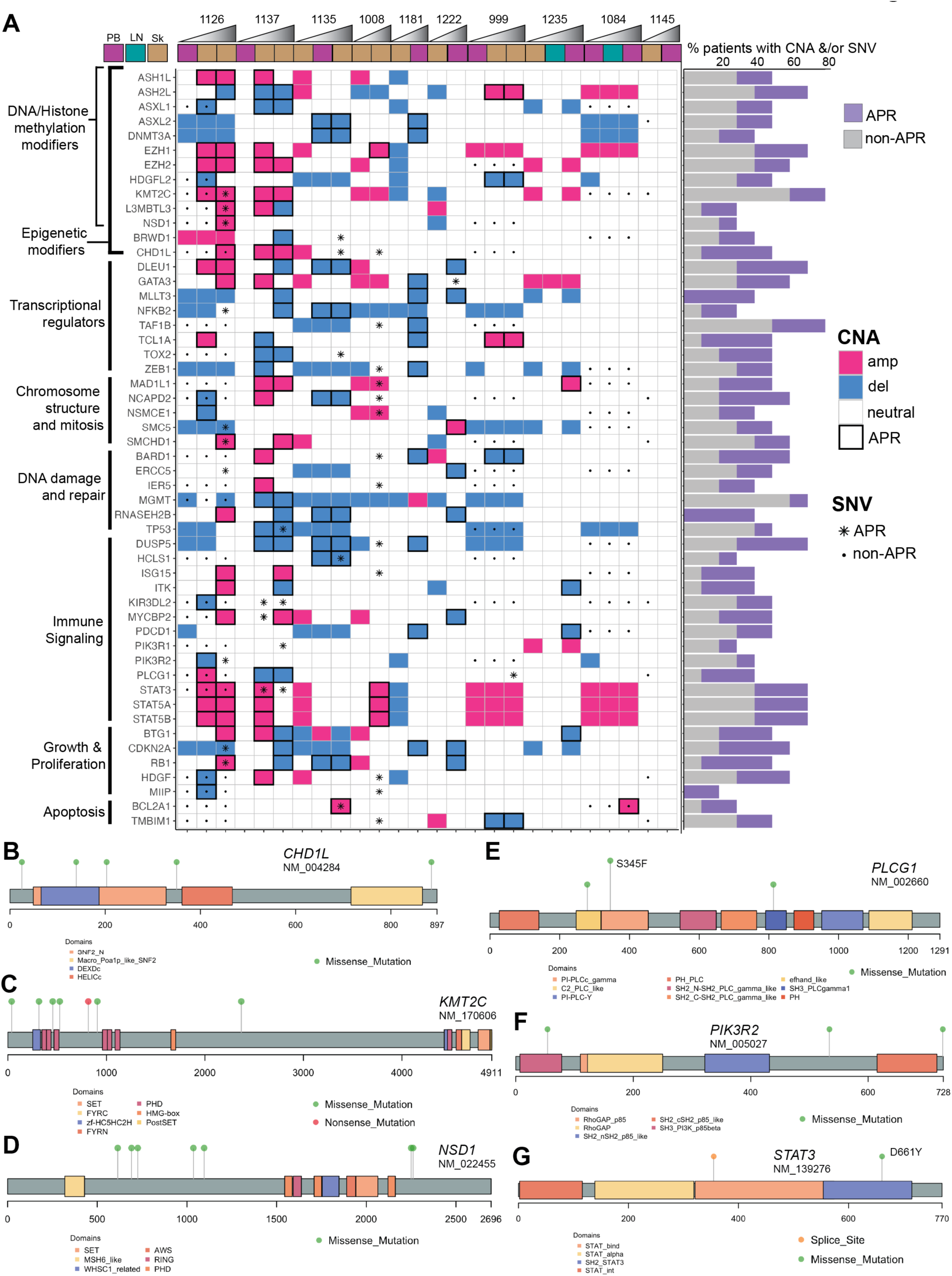
Epigenetic and transcriptional regulatory pathways are recurrently mutated with MF/SS disease and progression. **A)** Tile graph shows genome copy number amplifications (magenta) and deletions (blue) detected by WGS as well as somatic genome sequence variants detected by exome-seq (central black dots) for each gene (rows) and patient sample (columns). Specimen types are indicated by color blocks above the heatmap and are arranged in order from first to last (left to right) collected for each patient, time shown by triangles. APR CNA are indicated by thick black borders; APR sSNV are indicated by asterisks. In a histogram to the right of the heatmap, the percentage of patients with non-APR CNA and/or sSNV is indicated in grey, and that of APR CNA and/or sSNV is indicated in lavender. **B-G)** Lollipop diagrams show examples of six recurrently mutated proteins in the MF/SS cohort. Protein domains are indicated by shaded boxes, mutation locations and types are indicated by lollipops of different colors (missense, nonsense, splice site). (**B-D**) Mutations in epigenetic modifiers; (**E-G**) mutations in signaling proteins. Lollipop height is proportional to the number of samples with variants at the same amino acid residue. Except for S345F in the PI-PLCc_gamma domain of PLCG1, which is present in 2 samples, the rest of the mutations shown are present in one sample each.

### STAT3 mutation D661Y confers gain-of-function (GOF) activity

Given our finding of a D661Y somatic mutation in the SH2 protein interaction domain of the transcription factor STAT3, which is critical for activation and nuclear accumulation of STAT dimers^20^, we sought to define the functional impact of this amino acid change. To accomplish this, we assessed STAT3 transcriptional regulation in reporter assays and mapped genome-wide binding of STAT3 using CUT&RUN-sequencing. For these assays, we transfected wild type (WT) or D661Y mutant STAT3 plasmids into STAT3-null A4 cells^21^ and confirmed equal levels of expression (**Figures 4A, S4B**). Genomic regions bound by WT STAT3 or STAT3 D661Y were defined using a computational pipeline for peak calling and gene annotation similar to previous work^22^. **Figure 4B** shows that regions bound by WT STAT3 and STAT3 D661Y have similar proportional distributions among functional genomic categories (e.g., promoter, distal intergenic, intron). We next plotted WT STAT3- and STAT3 D661Y-bound regions within 3kb centered around transcription start sites (TSS) of genes (**Figure 4C**). This comparison showed a striking difference, with a greater number of STAT3 D661Y-bound regions overlying and within 3kb of TSS compared to WT STAT3. More than half of the peaks (4743, 62%) were common to both proteins, while 14% (1052) were unique to STAT3 and 25% (1902) were unique to STAT3 D661Y (**Figure 4D**), suggesting that the D661Y mutant gains more binding sites than it loses, which is consistent with **Figure 4C**. Pathway analysis showed that peaks common to both WT STAT3 and STAT3 D661Y include signaling by Rho GTPases and receptor tyrosine kinases, focal adhesion and FAK (focal adhesion kinase) pathways, which are involved in cell adhesion and motility, intra- and inter-cellular trafficking, and cell growth (**Figure 4D**). Pathways enriched in peaks unique to STAT3 D661Y include CDC42 (cell division control protein 42) GTPase cycle and several other Rho GTPase pathways, signaling by GPCR (G-protein coupled receptors), and diseases of signal transduction.

**Figure 4.**
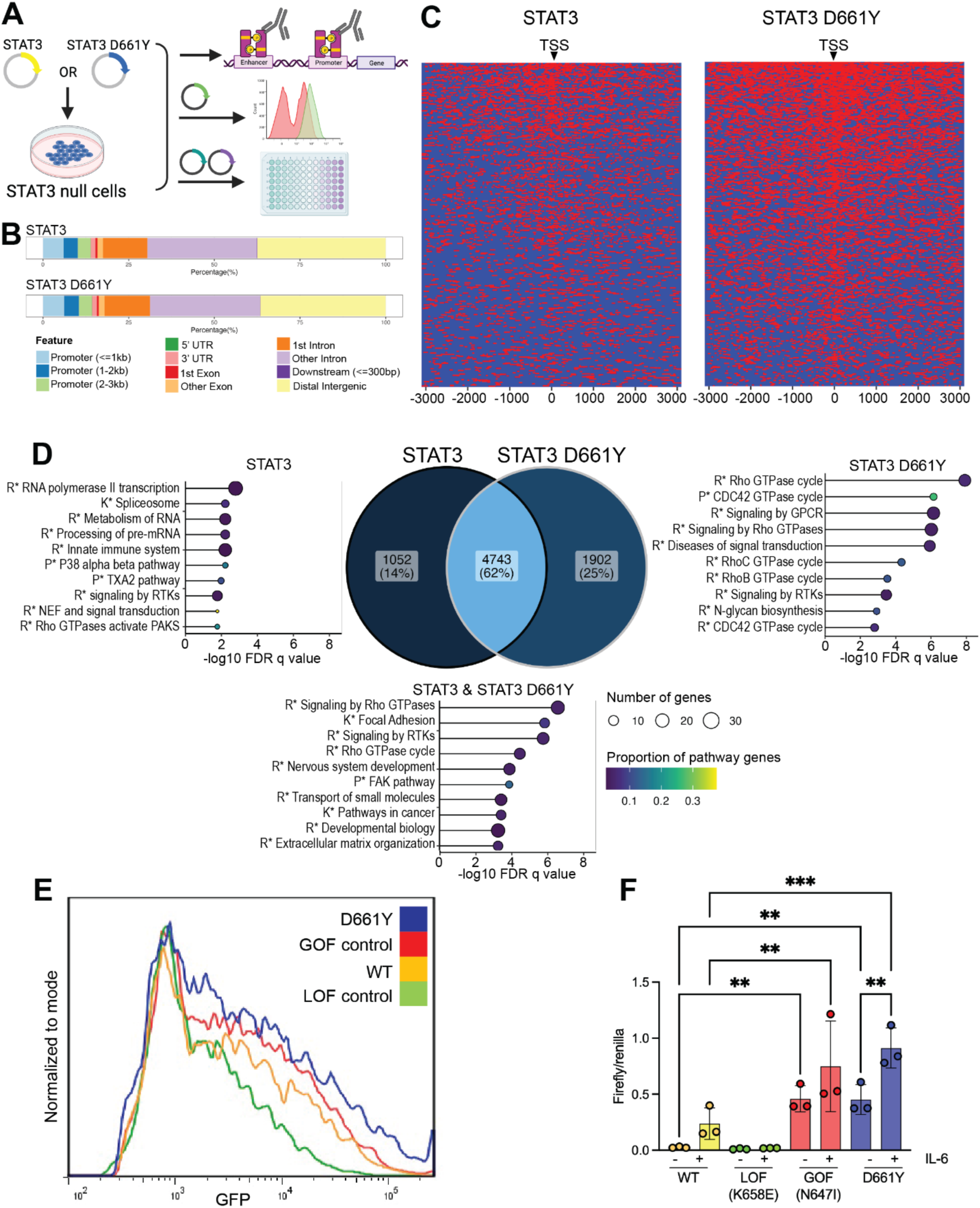
STAT3 D661Y is a gain of function mutation. **A)** STAT3 null A4 cells were transfected with plasmids encoding STAT3 or STAT3 D661Y then subjected to CUT&RUN-seq using a STAT3 antibody. For GFP-reporter assays, STAT3 null A4 cells were transduced with a STAT3-GFP reporter prior to transfection of STAT3, STAT3 D661Y, STAT3 LOF (K558E, loss of function), or STAT3 GOF (N647I, gain of function) plasmids. For luciferase assays, the same expression plasmids were transfected into STAT3 null A4 cells with STAT3-luciferase reporter and control renilla plasmids. **B)** Regions bound by STAT3 and STAT3 D661Y have similar proportional distributions among functional genomic categories. **C)** Heatmaps show increased genomic binding by STAT3 D661Y (right) compared to unmutated STAT3 (left) overlying and within 3kb of the transcription start site (TSS) of genes (CUT&RUNseq). **D)** Venn diagram shows the number of genes with binding of STAT3, STAT3 D661Y, or both within 3kb of the TSS, with D661Y having nearly twice as many unique genes bound compared to STAT3. Graphs show the top 10 pathways significantly enriched in each group of genes, as well as the number of genes found in each pathway (circles) and the proportion of the total genes in that pathway (color range) (CUT&RUNseq). **E)** Flow cytometry plot shows the number of cells normalized to mode and relative intensity of STAT3-driven GFP fluorescence for cells expressing STAT3, STAT3 D661Y, STAT3 LOF, or STAT3 GOF; STAT3 D661Y shows the highest intensity. **F)** Bar graph shows the STAT3-driven luciferase activity in cells expressing STAT3, STAT3 D661Y, STAT3 LOF, or STAT3 GOF at baseline and with IL-6 stimulation. ** p<0.01, *** p< 0.001 One-way ANOVA with Tukey’s multiple test correction. Additional significant comparisons not shown due to space constraints.

We next evaluated the impact of the D661Y mutation on the transcription of reporter genes. STAT3-null A4 cells were transfected with WT STAT3, STAT3 D661Y, and STAT3 gain or loss of function (GOF/LOF) mutant expression plasmids plus STAT3-GFP or STAT3-luciferase reporter plasmid. STAT3 D661Y had the highest intensity of GFP fluorescence, even higher than the STAT3 GOF control (**Figure 4E**). Untreated STAT3 D661Y has significantly higher baseline activity than untreated WT and in fact is similar to IL-6-treated WT STAT3. IL-6 treatment further increases STAT3 D661Y activity **(Figure 4F)**. Together, these data demonstrate that the D661Y substitution in STAT3 confers gain of function transcriptional activity *in vitro* and increases STAT3 binding near genes whose protein products play roles in cell adhesion, migration, and proliferation.

### scRNAseq identifies malignant T cell populations via TCR repertoire and expression profiling

We used single cell transcriptome plus T-cell receptor (TCR) V(D)J sequencing to generate expression and TCR repertoire profiles from 16 PBMCs samples from 6 CTCL patients (**Figure 1B**). Per sample, an average of 7,413 cells were sequenced, with a mean of 78,277 sequencing reads and a median of 1,693 unique genes detected per cell, comparable to previous studies^3,4^. Uniform manifold approximation and projection (UMAP^23^) was used to group cells by their expression profiles into 24 clusters (**Figures 5A-B, S2B**). Cell type annotation was performed as in^3,4^ (**Figures 5C, S5A-B**). Of 92,496 cells from all samples, the majority were CD4^+^ T cells (76,924) as expected due to the CD4+ enrichment we performed to enable detection of subclonal malignant T cell populations. The majority of CD4^+^ T cells had a central memory expression phenotype as expected from peripheral blood. Automated annotations were checked manually using canonical marker genes for T lymphocytes (*CD4*, *CD3D*, T cell receptor beta (*TRBC2*)) and CTCL (*TOX*, *KIR3DL2*, *PLS3*)^3,4,24–28^ (**Figure 5D**), which supports the cell type annotations. **Figure 5E** shows the top 10 differentially expressed genes for the largest 16 clusters. Genes in bold highlight selected markers for each cluster, including the aforementioned canonical CTCL genes. As expected, loss of CD7 was identified in several clusters.

**Figure 5.**
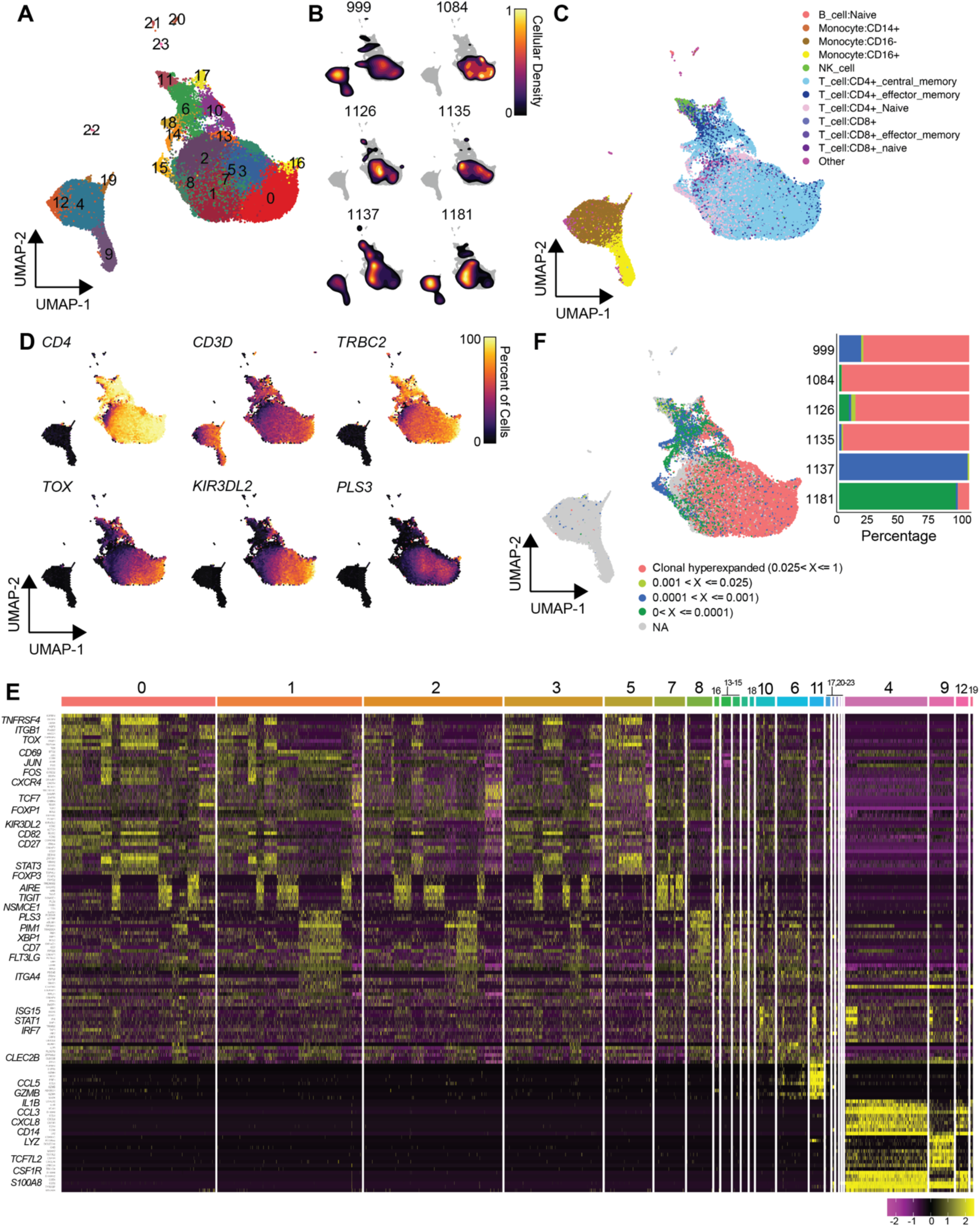
scRNAseq demonstrates hyperexpanded clonotypes and separation of benign and malignant T cells. **A)** UMAP projection of 92,496 cells from 16 peripheral blood samples from 6 CTCL patients across multiple time points identified 24 clusters of cells based on differential expression. **B)** Relative density of distribution for patient-specific cells along the UMAP. **C)** Cell assignments based on the highest spearman rho of purified immune populations in the Human Primary Cell Atlas (PMID: 24053356). **D)** UMAP projections depict percentages of total cells expressing selected canonical T-cell and CTCL genes. **E)** UMAP projection of the total cells overlaid with the clonal proportions of patient-specific TCR clonotypes. Stacked horizontal bar graph shows the percentage of repertoire space occupied by TCR clonotypes for each patient. **F)** Heatmap shows expression levels of the top 10 differentially expressed genes for each cluster. Selected gene symbols are highlighted in bold along the Y axis.

In T cell cancers, malignant T-cell populations express a single TCR clonotype^3,4,29^. We assigned the TCR clonotype for each cell using the scRepertoire (v1.3.2) R package previously developed by N.B.^30^. **Figure 5F** shows the UMAP projection of total cells overlaid with patient-specific TCR clonotypes colored by clonal proportion; hyperexpanded clonotypes were detected in 5 of 6 patients (**Figure S5C**). These clonotypes are expressed by cells with gene expression patterns consistent with malignant CTCL cells, including expression of canonical and established CTCL markers. In contrast, clonotypes with lower clonal proportions (<0.01) were expressed by cells with profiles consistent with non-malignant T cells (**Figure 5C-E**). In one patient (1137), there were no hyperexpanded TCR clonotypes detected, consistent with very low to no malignant T cells in the PBMC submitted for scRNA-seq. Skin samples subjected to eWGS from this patient demonstrated genomic CNA consistent with CTCL; most of these changes were not detected in the single PBMC eWGS sample (**Figure 2A**).

Using the TCR clonotypes and gene expression-based cell annotations, we defined three groups of cells: malignant CD4+ T cells, non-malignant CD4 or CD8 T cells, and non-T cells (e.g., monocytes, B cells, NK cells) (**Figure S5D**). Many of the same genes identified in the full clustering analysis (**Figure 5E**) were significantly different in comparing these three groups, including canonical CTCL, cell adhesion, and signaling gene pathways (**Figure S5E**). In summary, scRNA-seq of T-cell enriched CTCL samples identified cells with expression patterns and TCR clonotype skewing consistent with CTCL cells and segregated these from non-malignant T and non-T cells. Notably, these expression profiles mirror patterns of disease progression and epigenetic regulation that we previously identified, including for T-cell regulation (*AIRE*, *FOXP3*, *KIR3DL2*, *TIGIT*), T-cell transcription factors (*JUN*, *FOS*, *TCF7*), cell adhesion, chemotaxis, signaling, and apoptotic pathways that we previously identified using epigenomic approaches, including *CXCR4*, *ITGB1*, and *BCL2*^2^.

### scRNAseq detects clonal evolution of CNA and sSNV

Having identified CNA, sSNV, and expression changes in bulk sequencing studies, we next sought to define clonal evolution at single cell resolution using scRNAseq. We highlight two patients as examples of distinct temporal patterns (**Figure 6A-B**). UMAP projections and bar graphs in **Figures 6C-E, S6A-F** show the numbers and proportions of single cells by TCR clonotype, cell annotation, and cell type (malignant T, non-malignant T, and non-T) as defined in **Figure S5**. Samples from patients 1084 and 1126 demonstrate substantial enrichment of CTCL cells (>70%), while samples from patient 1181 contains fewer CTCL cells (**Figures 6C-E, S6A-F)**. Consistent with these findings, clinical records indicated that patient 1181 had a low burden of peripheral blood disease at these time points.

**Figure 6.**
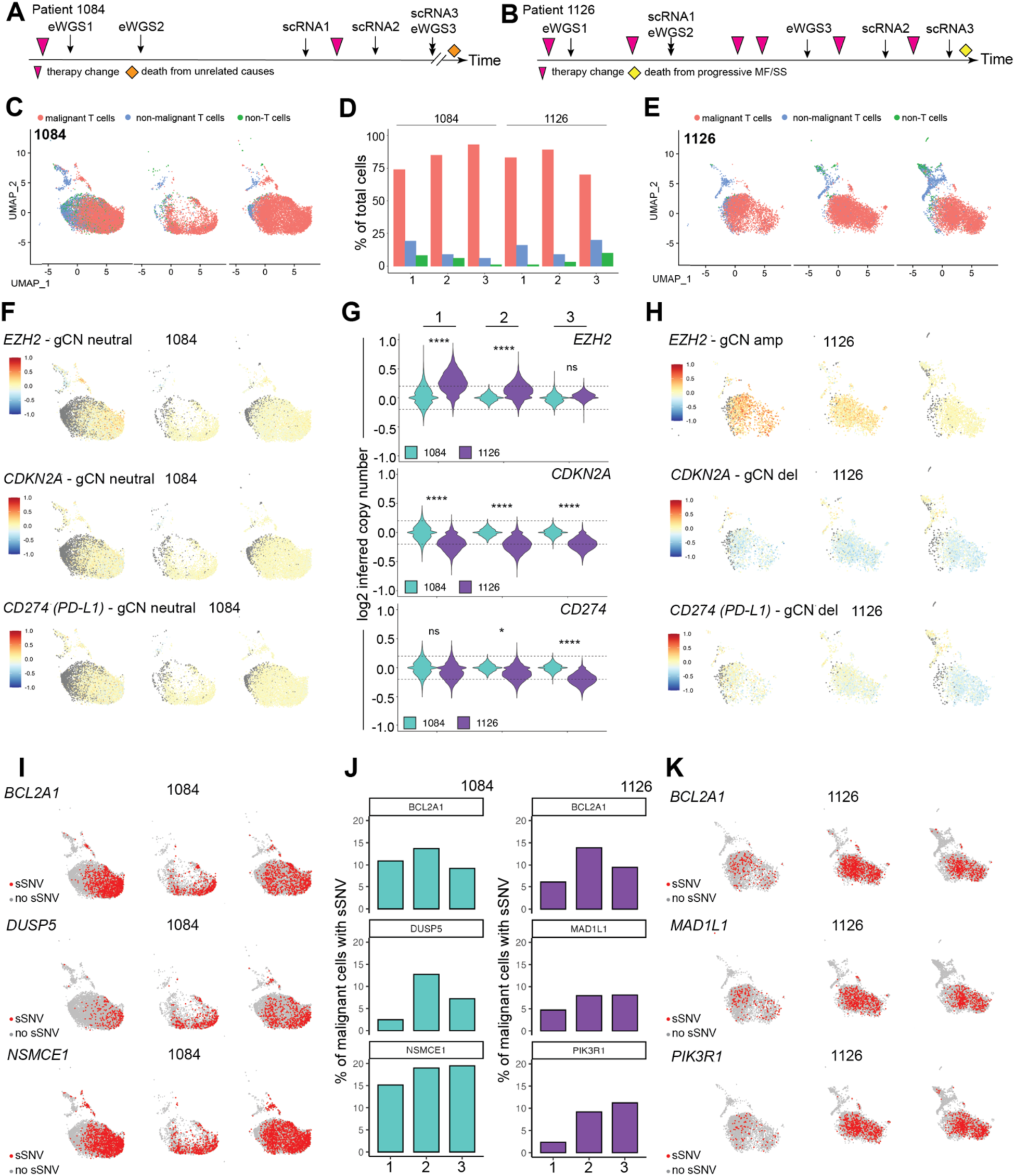
Genome copy number alterations and single nucleotide variants are detectable in scRNAseq and vary with therapy response and disease progression. A-B) Timelines depicting relative times for sequencing and changes in therapy for patients 1084 and 1126 are shown. UMAP projections of individual cells from 3 serial specimens per patient from patients 1084 (**C,F,I**) and 1126 (**E,H,K**); time from first specimen increases from left to right and corresponds to the scRNAseq points on the timelines **(A-B)**. Cells assigned to malignant T (red), non-malignant T (blue), or non-T (green) cell annotations depicted by UMAP **(C,E)** and percent bar plot **(D)**. Inferred copy number (iCN) for individual cells as determined from scRNAseq data and shown by UMAP (**F,H**) and violin plot **(G)** for selected genes in epigenetic modifier, cell cycle, and transcription factor pathways that have diploid (abs[log2 iCN] < 0.2) or altered (abs[log2 iCN] >0.2) iCN in serial samples is depicted. WGS initially identified CNA (gCN = genomic copy number determined by WGS). (**I-K**) UMAPs depict sSNV in selected genes in apoptosis, DNA repair, and signaling pathways detected in scRNAseq data **(I,K)**; percentages of cells with sSNV are shown by bar plot **(J)**. sSNV were initially identified in exome sequencing and met all criteria as stated in Methods.

We next evaluated CNA in scRNAseq by first calculating single-cell inferred copy number estimates. **Figure 6F-H** and **S6G-J** depict examples of genes identified in a CNA (defined by matched WGS) for one patient, while the other patient had neutral (normal diploid) copy number. These examples show heterogeneity of gene copy number across TCR clonal CTCL cells, demonstrating subclonal populations with the CNA and some without. Higher inferred copy number for *EZH2* was detected in a subset of malignant T cells in scRNAseq samples 1 and 2 from patient 1126. WGS performed on samples collected prior to or simultaneously with these two samples also detected amplification of *EZH2* (**Figures 2, 6B**). The number of cells with amplification of *EZH2* decreased over time (63% to 28% to 0.8%) (**Figure 6G-H)**. In contrast, the proportion of cells with lower inferred copy number of *CDKN2A* (1126) remains relatively constant (47%-53%-49%) (**Figure 6G-H)**; a similar pattern is seen for *NFKB2*, *DNMT3A*, and *ZEB1* **(Figure S6G-I)**. In yet a different pattern, the proportion of cells with deletion of *CD274* (PD-L1) increases over time (17%-19.5%-49%) (1126, **Figures 6G-H, S6J**). CNA deletion and reduced inferred copy number of *CD274* and *PDCD1* were also detected in 1181, but *PDCD1* copy number was normal in 1126 (**Figure S6J-K**). These changes may indicate shifts in subclonal populations due to differences in therapy response and/or disease progression for patient 1126, whose disease progressed on multiple therapies and who ultimately died of CTCL seven months after the last scRNAseq sample was collected (**Figure 6B**). In contrast, patient 1084 had relatively stable disease, exhibiting few acquired molecular alternations and requiring few therapeutic changes; the patient ultimately died of unrelated causes two years after the last sample collection.

We next performed global comparisons of CNA and single cell gene expression. We found that 3,777 genes were differentially expressed in CTCL cells compared to non-malignant T and non-T cells (padj<0.05 & abs(average log2 fold change) >0.5); of these, 2,681 had concordant changes in genome copy number determined by WGS (71% of differentially expressed genes). These include several of the genes with recurrent CNA and sSNV (*GATA3*, *NSMCE1*, *TMBIM1*, *MAD1L1*, *ASH2L*, *ASXL2*, *NSD1,* **Figure 3A**). Four additional genes’ correlative expression was also detected by bulk RNAseq (*EZH2*, *STAT3*, *TP53*, *PIK3R1*; **Figures 2, S3**). Thus, single cell mapping of malignant CTCL cells further supports a link between genomic CNA, the most common type of genomic mutation in CTCL, and disrupted gene expression in individual malignant CTCL cells.

We next sought to evaluate clonal evolution of sSNV at single cell resolution. To map sSNV to individual cells, we generated patient-specific lists of sSNV based on exome sequencing results and identified the cell-barcoded sequencing reads that contained these variant sequences. We further set filters for barcode frequency and required detection in CTCL cells but not in non-malignant T or non-T cells. **Figure 6I-K** shows examples of sSNV that meet these criteria: UMAPs for three serial samples from two patients depict cells in red that contain exome seq-defined sSNV in the indicated gene. Like the CNA shown in **Figure 6F-H**, these examples also show heterogeneity of sSNV detection within TCR clonal cells, demonstrating subclonal populations of malignant T cells with or without the sSNV. *BCL2A1*, *DUSP5*, *MAD1L1*, *NSMCE1*, and *PIK3R1* variants were detected in a subset of malignant clonal T cells in patients 1084 and 1126, compare **Figure 6I** to **6C** and **6K** to **6E**. In addition, these plots demonstrate that the proportions of cells with sSNV generally show a more stable pattern for 1084 compared to 1126 (**Figure 6I-K**, quantified in **6J**). These changes in the prevalence of cells expressing sSNVs also may reflect distinct subclonal populations with differences in response to therapy and/or disease progression.

### Genomic alterations associate with response to therapy

We next evaluated molecular changes identified in this study for their association with response to specific therapies (**Table 1**), first focusing on therapeutic targets. Seven multi-omics cohort patients received mogamulizumab, whose target is CCR4; 2 of 3 non-responders had APR deletion of *CCR4* while 1 of 4 responders had non-APR *CCR4* deletion at a single timepoint. Five multi-omics cohort patients received checkpoint inhibitors targeting PD-1/PD-L1; 1 of 2 non-responders had non-APR deletions of *PDCD1* (PD-1) at a single timepoint and *CD274* (PD-L1) at all timepoints; no responders had alterations in these genes. Three multi-omics cohort patients received PI3K inhibitors; all had APR deletions and/or sSNV in PI3K regulators (PTEN, PIK3R1 and PIK3R2) and none responded (1 of 3 patients without eWGS responded). There was no significant difference in gene expression between response and resistance groups for these genes (**Figure S7A-F**). These results demonstrate that mutation of therapeutic targets, particularly when acquired with progression or detected at multiple timepoints, is associated with lack of therapeutic response.

Because HDACi inhibit several histone deacetylases that themselves have many downstream targets^2,31–35^, we performed unbiased evaluation of all genes with CNA and/or sSNV for association with HDACi response using Fisher’s exact test. We identified genomic alterations in several genes not previously associated with response to HDACi (p < 0.05), including genes clustered within a deleted region in chromosome 3p24.1 (*EOMES*) and an amplified region within 15q26.1 (e.g., *BLM*, *IDH2, IQGAP1*) (**Figure S7G-H**). Gene expression was not significantly different between the drug response groups. Taken together, these results may indicate an association of these genomic alterations with response to HDACi, though larger studies are needed to confirm these findings.

## DISCUSSION

CTCL remains a challenging disease due to its significant heterogeneity, therapy resistance, and relentless progression. Here we present a comprehensive multi-omics view of CTCL evolution, leveraging clinically annotated serial samples to dissect the molecular underpinnings of CTCL progression with the goal of identifying clinically useful biomarkers and therapeutic targets.

We identified several recurrent APR mutation patterns. Similar to previous reports^36–40^, we found that mutation of therapeutic targets (CCR4, PD-1, PD-L1) and therapy-associated factors (PI3K regulators PTEN, PIK3R1 and PIK3R2) was associated with lack of response to therapies targeting these factors or pathways, particularly when APR and/or at multiple timepoints. We also identified mutations associated with HDACi response/resistance, including genes in Rho GTPase pathways, which we previously reported to have activated chromatin and increased expression in HDACi resistance^2^. Particularly remarkable is *IQGAP1*, which we showed is a specific target of D661Y mutant STAT3 and is significantly overexpressed in CTCL. IQGAP1 is a scaffold protein that assembles multiple protein signaling complexes, including PI3K/Akt, MAPK, STAT, and GTPases, which control cytoskeleton organization, cell migration, and cell proliferation^41–45^. Consistent with previous reports using single CTCL samples^6,14,46,47^, we found APR mutations and overexpression of *STAT3*. We also identified a gain of function sSNV in the SH2 domain of STAT3 (D661Y) that caused enhanced binding to genes in Rho GTPase pathways. Together, these findings demonstrate that CTCL progression is associated with genomic changes in therapeutic targets and provide further support for a previously unrecognized role for Rho GTPase pathway dysregulation in CTCL pathogenesis.

Our studies also revealed progression-associated changes in epigenetic pathways, most strikingly in EZH2, the catalytic subunit of the polycomb repressive complex 2 (PRC2). Genetic alterations and overexpression of EZH2 are common in B-cell lymphomas, consistent with its role in regulating B lymphocyte differentiation^48–55^. Notably, non-canonical functions of EZH2 are also oncogenic, including interactions with DNA methyltransferases and methylation of transcription factors, such as STAT3^56,57,57–63^. EZH1/2 dual inhibitors have shown efficacy in B and T cell malignancies with or without EZH2 mutations^52,64–69^. Intriguingly, combination EZH2 and HDACi, the latter of which is well-established in CTCL treatment^2,35,70,71^, showed promise in aggressive prostate cancer models^72^. These data suggest that EZH1/2 inhibition may also benefit patients with CTCL.

Taken together, our results present a comprehensive multi-omics view of clonal evolution in CTCL and reveal molecular changes that are frequently acquired with disease progression and therapy resistance. Our findings support an approach in which genomic analysis is widely utilized for improved disease monitoring, biomarker-informed clinical trial design, and genome-guided therapeutic decision making. Moreover, these molecular changes present new opportunities for therapeutic targeting; indeed, we have initiated a Phase I study of EZH2 inhibition in CTCL patients.

## Acknowledgements

This work was supported by an American Heart Association Predoctoral Fellowship grant #P23-01044 to HKD; Washington University BioSURF funding to ECM; NIH P01AI155393 and CDI Center for Pediatric Immunology at Washington University and St. Louis Children’s Hospital to MAC; the Siteman Cancer Center Foundation, the Barnes-Jewish Hospital Foundation Steinbeck Designated Fund, and McDonnell Genome Institute and Illumina research support to NMS and JEP. Sequencing studies performed at the McDonnell Genome Institute Genome Technology Access Center were partially supported by NCI Cancer Center Support Grant #P30 CA91842 to the Siteman Cancer Center, the National Center for Research Resources (NCRR), and the Institute of Clinical and Translational Sciences (ICTS) at Washington University in St. Louis, which is funded by the National Institutes of Health’s NCATS Clinical and Translational Science Award (CTSA) program grant #UL1 TR002345. We also acknowledge the Research Infrastructure Services (RIS) group at Washington University in St. Louis for providing computational resources and services. We thank Marcus Watkins for helpful comments on the manuscript. We are grateful to the clinical coordinators, the patients and their families without whom this study would not have been possible.

## Authorship Contributions

JEP and NMS designed the study. NMA and AM recruited patients for the study. HKD, JMA, CCQ, JAS, MTH, NCB, JA, MW, AF, and JEP coordinated and processed clinical samples, performed experiments, analyzed data, and generated figures. HKD, JMA, NCB, NMS and JEP wrote the manuscript with input from all other authors.

---

**Figure S1.**
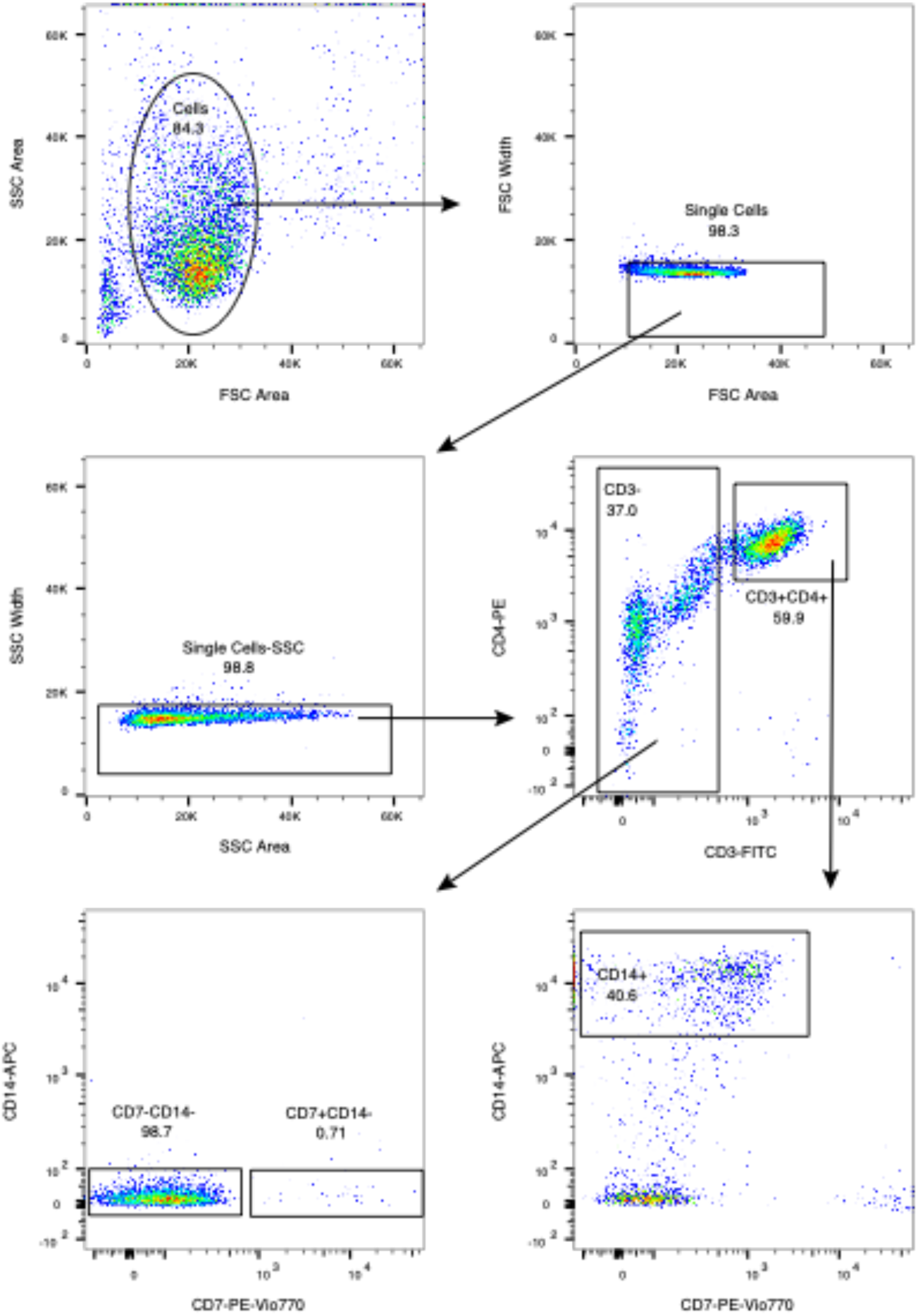
Example plot of flow cytometry gating. PBMC from a patient were isolated, incubated with fluorescence-conjugated antibodies as shown, and subjected to flow cytometry or FACS with gating as shown.

**Figure S2.**
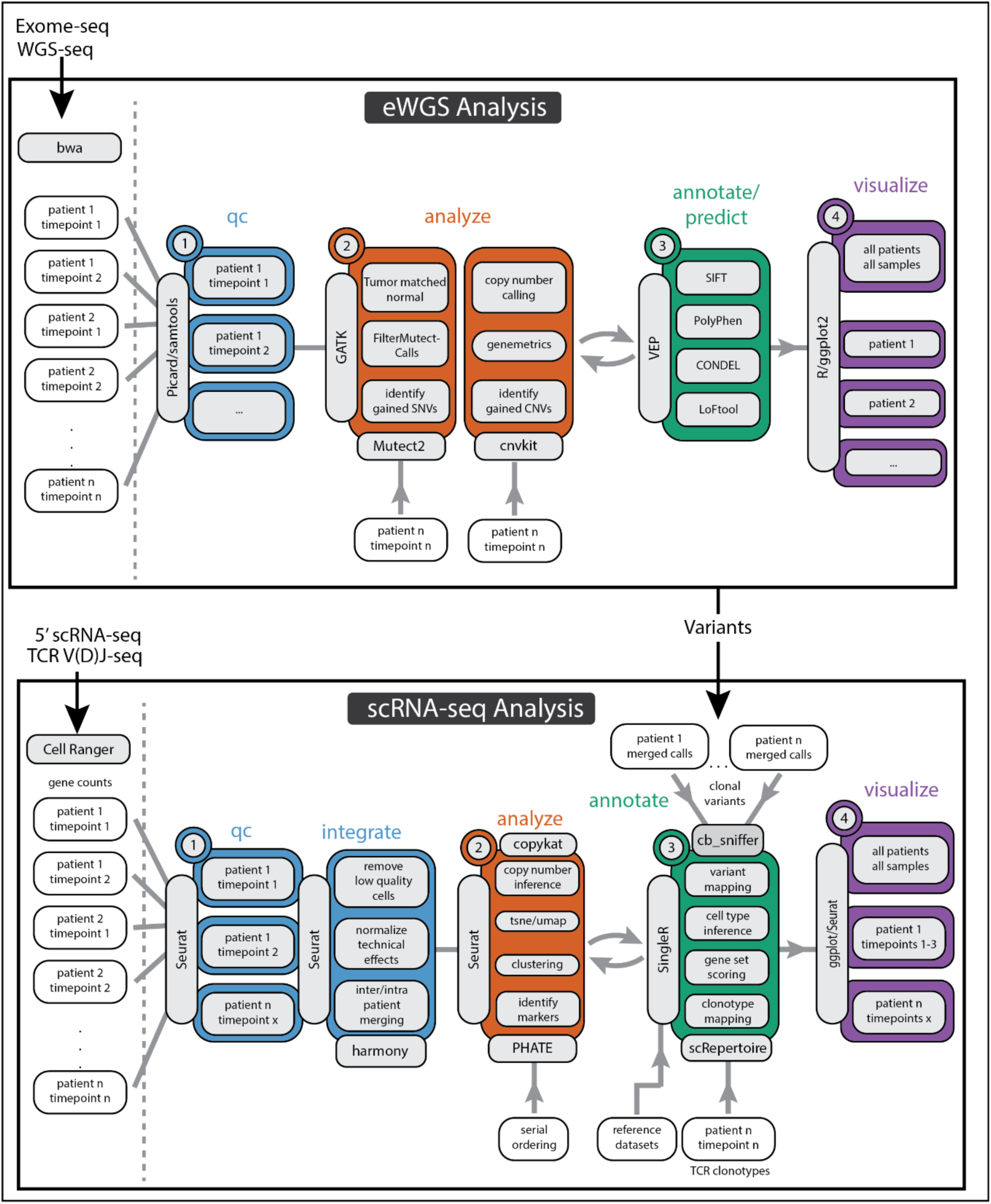
Analysis pipelines for exome, whole genome, and single cell RNA sequencing. Data flow and software programs are shown. See Methods for details.

**Figure S3.**
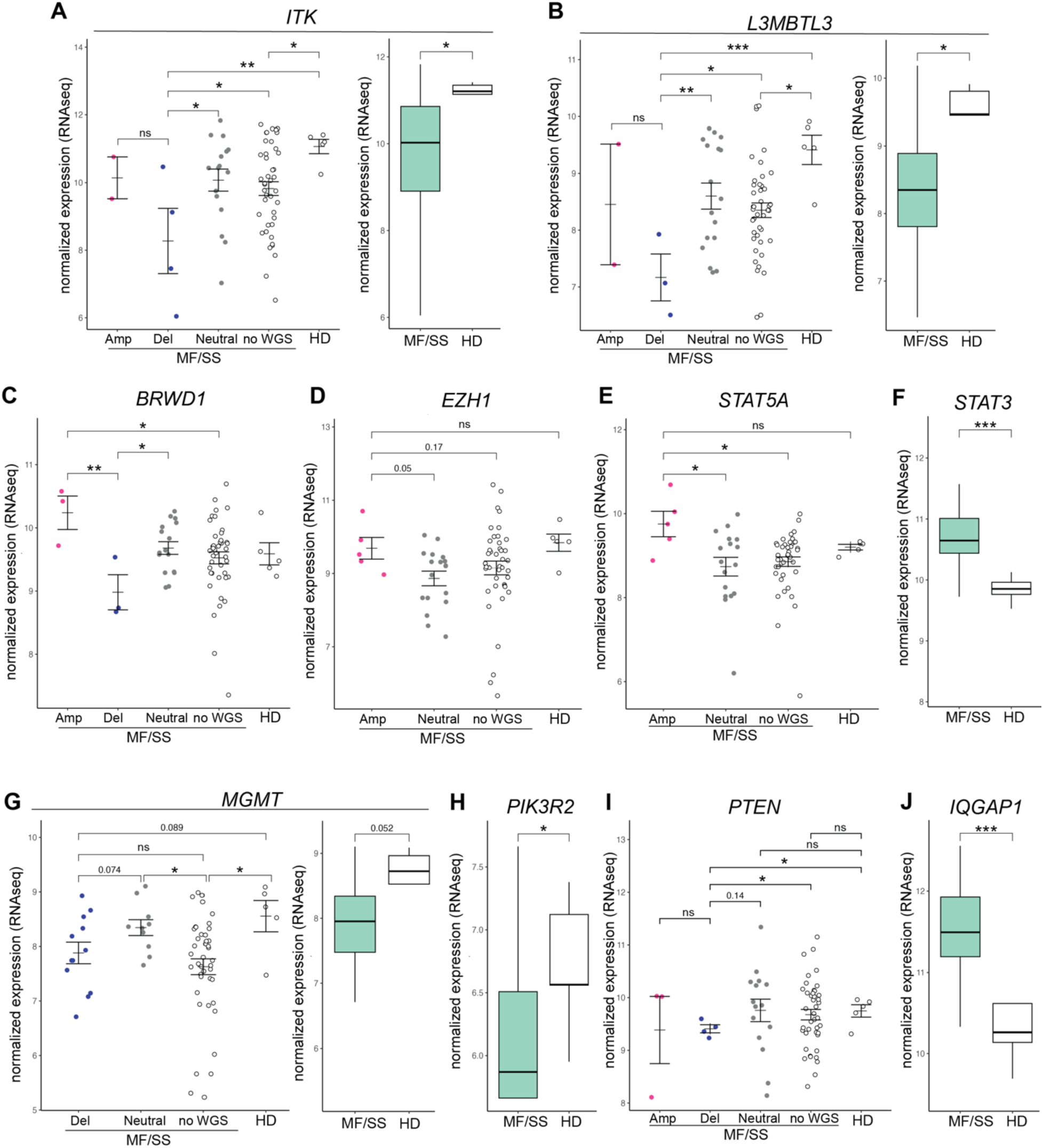
Copy number alterations correlate with gene expression and disease status. Grouped dot plots depict normalized expression (bulk RNAseq) of example genes within CNA; CNA groups are based on WGS performed at the closest point in time to the RNAseq. Box plots depict normalized expression (bulk RNAseq) of example genes with samples separated by disease status. HD – healthy donor. Wilcoxon rank sum test * p<0.05, ** p<0.01, *** p<0.001.

**Figure S4.**
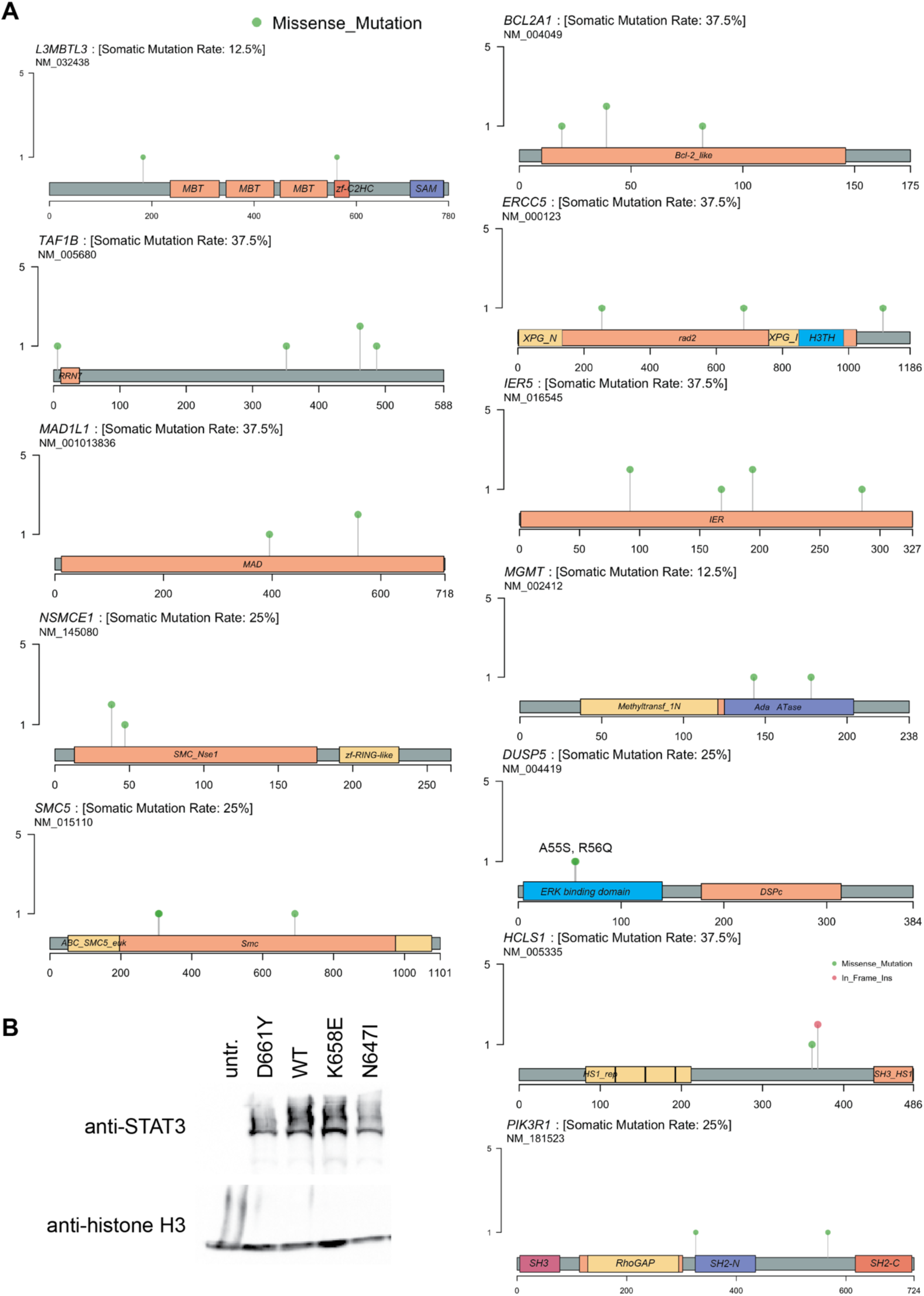
Epigenetic and transcriptional regulatory pathways are recurrently mutated with MF/SS disease and progression. **A)** Lollipop diagrams show examples of recurrently mutated proteins in the MF/SS cohort. Protein domains are indicated by shaded boxes, mutation locations and types are indicated by lollipops of different colors (missense, nonsense, splice site). Lollipop height is proportional to the number of patients (may be multiple samples per patient) with variants at the same amino acid residue. The sSNVs in *DUSP5* in two patients were located at adjacent amino acids (A55S, R56Q) in the ERK binding domain. **B)** Western blot of A4 STAT3 null cells untransfected (untr.) or transfected with the indicated *STAT3* expression plasmids shows equal expression of all STAT3 variants.

**Figure S5.**
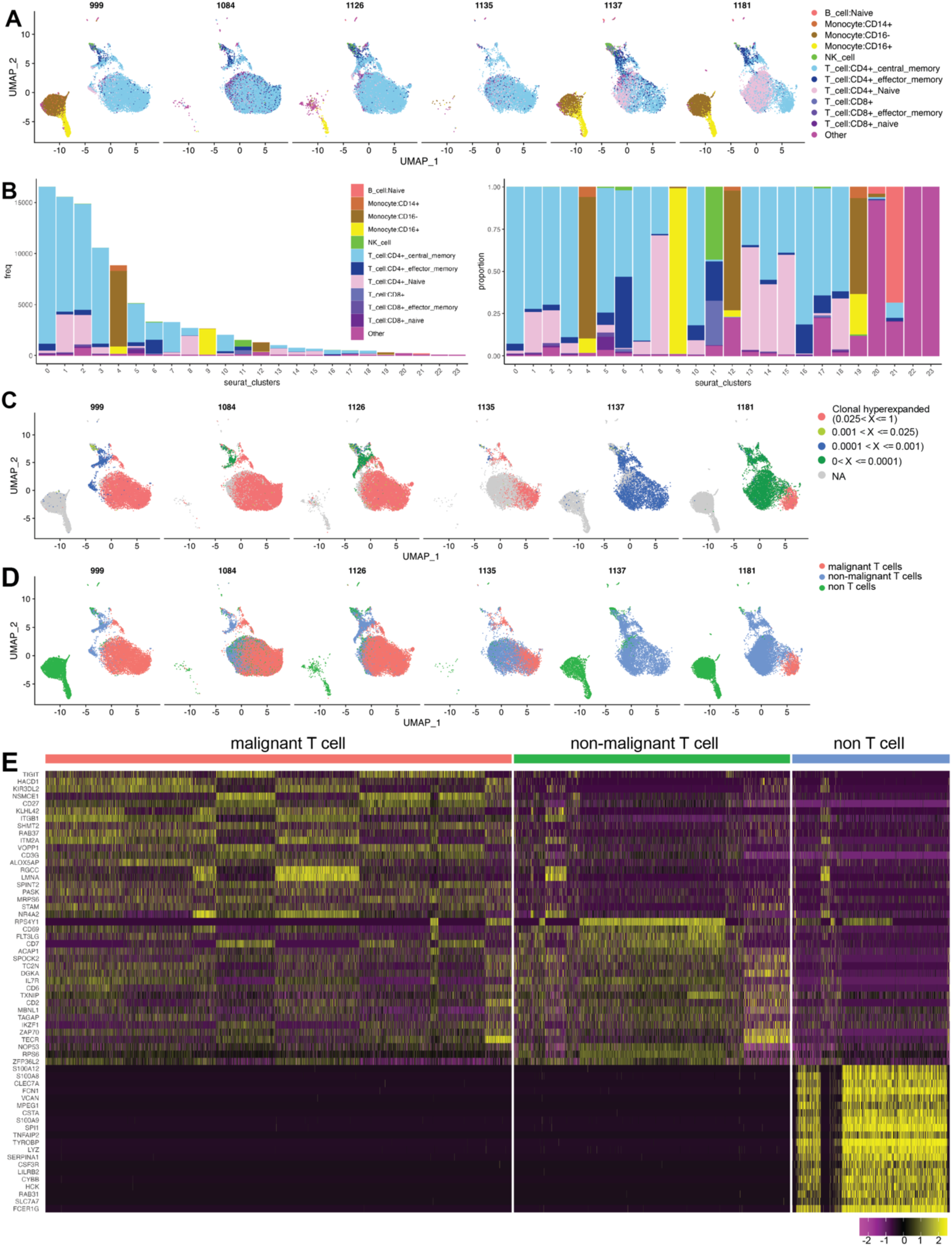
scRNAseq demonstrates hyperexpanded clonotypes and separation of benign and malignant T cells. UMAP projections and stacked horizontal bar graphs show the number and proportion of cells by annotation groups **(A-B),** relative abundances of TCR clonotypes **(C),** and malignant T, non-malignant T, and non-T cells **(D)** for scRNAseq samples from each of 6 patients. **D)** Heatmap shows expression levels of the top 10 differentially expressed genes ranked by log2 fold change for each category of malignant T, non-malignant T, and non-T cells.

**Figure S6.**
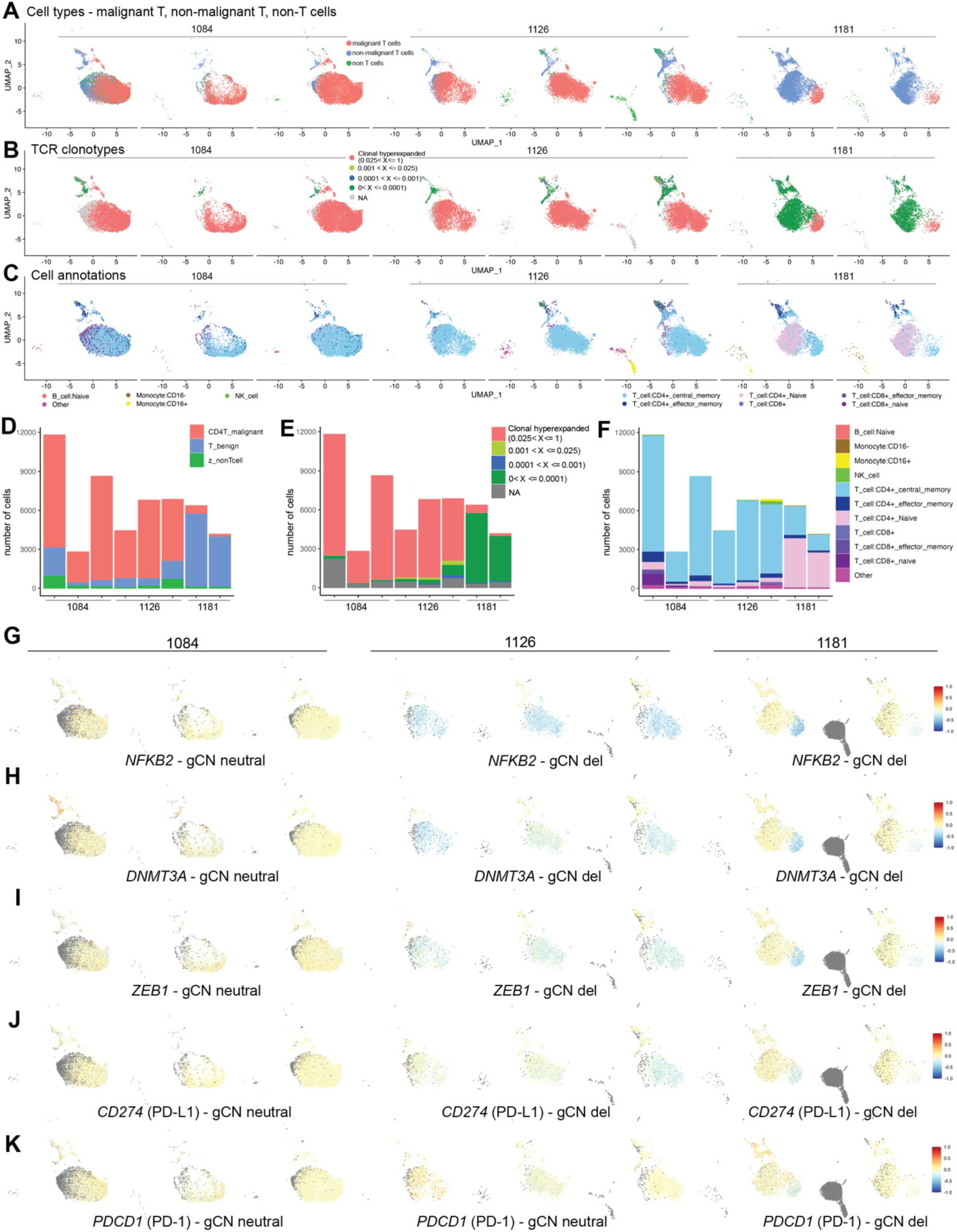
Genome copy number alterations are detectable in scRNAseq and vary with therapy response and disease progression. (**A-C**) UMAP projections of individual cells from 8 individual serial specimens from 3 patients; time from first specimen increases from left to right for each patient. Cells are colored by groups, cell type (**A**), TCR clonotype (**B**), and annotation (**C**), as defined in **Figure S5 A-D**. (**D-E)** Stacked bar graphs show the number of cells of each cell type (**D**), TCR clonotype (**E**), and cell annotation (**F**) for each sample (as in **A** - **C**). (**G – K**) UMAP projections show genomic copy number inferred from scRNAseq data of individual cells for 5 genes with genome copy number (gCN) alterations as indicated.

**Figure S7.**
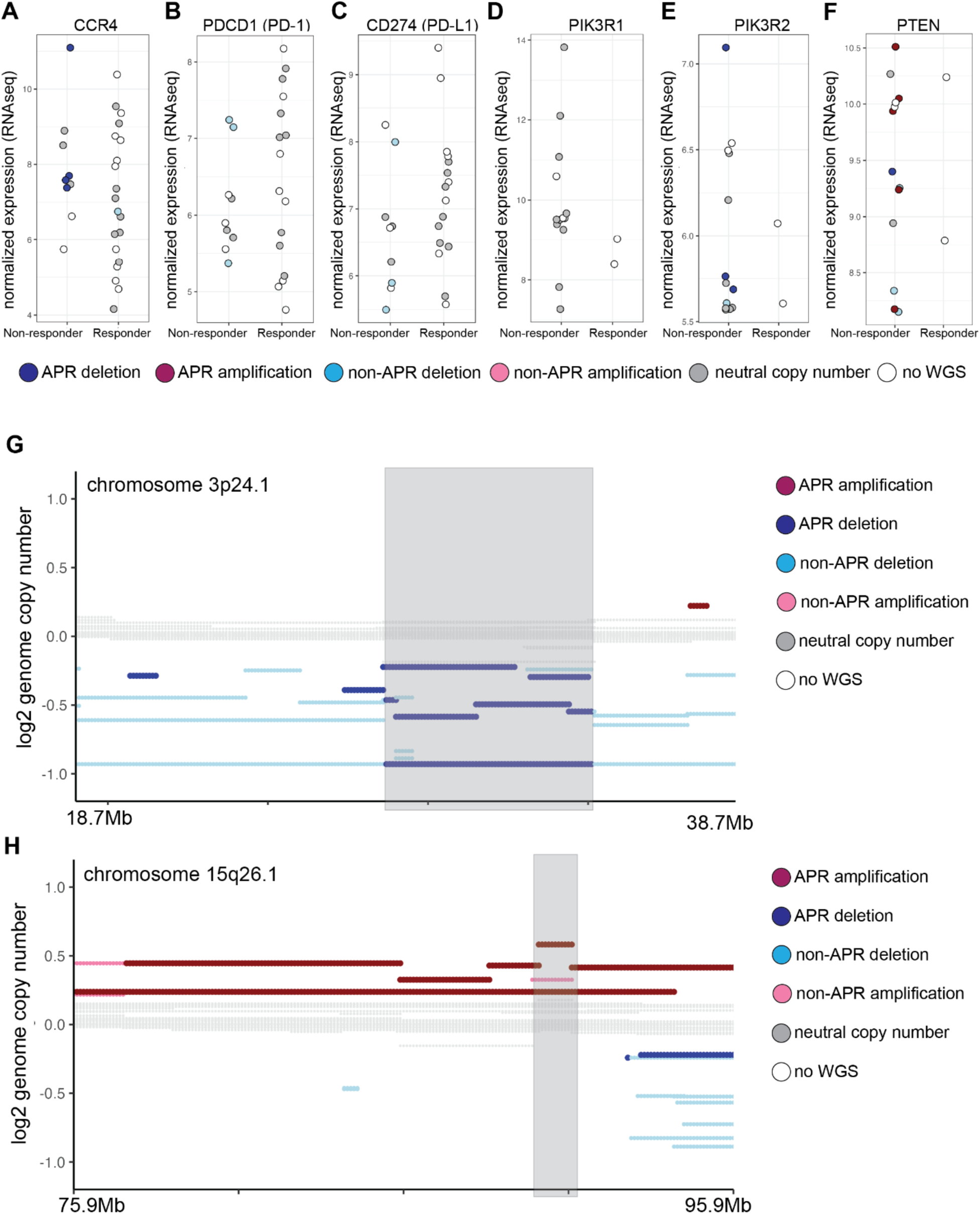
Genomic alterations associate with response to therapy. **(A-F)** Grouped dot plots depict normalized expression (bulk RNAseq) from patients grouped by response or not to mogamulizumab (**A**), checkpoint inhibition (**B-C**), or PI3K inhibition (**D-F**). CNA detected at any timepoint is indicated by color. Some patients have multiple RNAseq samples depicted. (**G-H**) Genome copy number plots depict log2 genome copy number for two regions associated with response to HDACi.

## REFERENCES

1. Mehta-Shah N, Horwitz SM, Ansell S, et al. NCCN Guidelines Insights: Primary Cutaneous Lymphomas, Version 2.2020. J Natl Compr Canc Netw. 2020;18(5):522–536.

2. Andrews JM, Schmidt JA, Carson KR, et al. Novel cell adhesion/migration pathways are predictive markers of HDAC inhibitor resistance in cutaneous T cell lymphoma. EBioMedicine. 2019;

3. Borcherding N, Severson KJ, Henderson NT, et al. Single-cell analysis of Sézary syndrome reveals novel markers and shifting gene profiles associated with treatment. Blood Adv. 2022;bloodadvances.2021005991.

4. Borcherding N, Voigt AP, Liu V, et al. Single-Cell Profiling of Cutaneous T-Cell Lymphoma Reveals Underlying Heterogeneity Associated with Disease Progression. Clin Cancer Res. 2019;25(10):2996–3005.

5. Buus TB, Willerslev-Olsen A, Fredholm S, et al. Single-cell heterogeneity in Sézary syndrome. Blood Advances. 2018;2(16):2115–2126.

6. Choi J, Goh G, Walradt T, et al. Genomic landscape of cutaneous T cell lymphoma. Nat. Genet. 2015;47(9):1011–1019.

7. da Silva Almeida AC, Abate F, Khiabanian H, et al. The mutational landscape of cutaneous T cell lymphoma and Sézary syndrome. Nat. Genet. 2015;47(12):1465–1470.

8. Litvinov IV, Tetzlaff MT, Thibault P, et al. Gene expression analysis in Cutaneous T-Cell Lymphomas (CTCL) highlights disease heterogeneity and potential diagnostic and prognostic indicators. Oncoimmunology. 2017;6(5):e1306618.

9. McGirt LY, Jia P, Baerenwald DA, et al. Whole-genome sequencing reveals oncogenic mutations in mycosis fungoides. Blood. 2015;126(4):508–519.

10. Park J, Daniels J, Wartewig T, et al. Integrated Genomic Analyses of Cutaneous T Cell Lymphomas Reveal the Molecular Bases for Disease Heterogeneity. Blood. 2021;blood.2020009655.

11. Park J, Yang J, Wenzel AT, et al. Genomic analysis of 220 CTCLs identifies a novel recurrent gain-of-function alteration in RLTPR (p.Q575E). Blood. 2017;130(12):1430–1440.

12. Qu K, Zaba LC, Satpathy AT, et al. Chromatin Accessibility Landscape of Cutaneous T Cell Lymphoma and Dynamic Response to HDAC Inhibitors. Cancer Cell. 2017;32(1):27–41.e4.

13. Song X, Chang S, Seminario-Vidal L, et al. Genomic and single-cell landscape reveals novel drivers and therapeutic vulnerabilities of transformed cutaneous T-cell lymphoma. Cancer Discovery. 2022;candisc.1207.2021.

14. Wang L, Ni X, Covington KR, et al. Genomic profiling of Sézary syndrome identifies alterations of key T cell signaling and differentiation genes. Nat. Genet. 2015;47(12):1426–1434.

15. Christopher MJ, Petti AA, Rettig MP, et al. Immune Escape of Relapsed AML Cells after Allogeneic Transplantation. N Engl J Med. 2018;379(24):2330–2341.

16. Petti AA, Khan SM, Xu Z, et al. Genetic and Transcriptional Contributions to Relapse in Normal Karyotype Acute Myeloid Leukemia. Blood Cancer Discov. 2022;3(1):32–49.

17. Petti AA, Williams SR, Miller CA, et al. A general approach for detecting expressed mutations in AML cells using single cell RNA-sequencing. Nat Commun. 2019;10(1):3660.

18. Masle-Farquhar E, Jackson KJL, Peters TJ, et al. STAT3 gain-of-function mutations connect leukemia with autoimmune disease by pathological NKG2Dhi CD8+ T cell dysregulation and accumulation. Immunity. 2022;55(12):2386–2404.e8.

19. Vogel TP, Milner JD, Cooper MA. The Ying and Yang of STAT3 in Human Disease. J Clin Immunol. 2015;35(7):615–623.

20. de Araujo ED, Orlova A, Neubauer HA, et al. Structural Implications of STAT3 and STAT5 SH2 Domain Mutations. Cancers (Basel). 2019;11(11):1757.

21. Yang J, Huang J, Dasgupta M, et al. Reversible methylation of promoter-bound STAT3 by histone-modifying enzymes. Proc Natl Acad Sci U S A. 2010;107(50):21499–21504.

22. Cubitt CC, Wong P, Dorando HK, et al. Induced CD8α identifies human NK cells with enhanced proliferative fitness and modulates NK cell activation. J Clin Invest. 2024;e173602.

23. McInnes L, Healy J, Melville J. UMAP: Uniform Manifold Approximation and Projection for Dimension Reduction. 2020;

24. Litvinov IV, Netchiporouk E, Cordeiro B, et al. The Use of Transcriptional Profiling to Improve Personalized Diagnosis and Management of Cutaneous T-cell Lymphoma (CTCL). Clin Cancer Res. 2015;21(12):2820–2829.

25. Roelens M, de Masson A, Ram-Wolff C, et al. Revisiting the initial diagnosis and blood staging of mycosis fungoides and Sézary syndrome with the KIR3DL2 marker. Br J Dermatol. 2020;182(6):1415–1422.

26. Tang N, Gibson H, Germeroth T, et al. T-plastin (PLS3) gene expression differentiates Sézary syndrome from mycosis fungoides and inflammatory skin diseases and can serve as a biomarker to monitor disease progression. Br J Dermatol. 2010;162(2):463–466.

27. Dulmage BO, Akilov O, Vu JR, Jr LDF, Geskin LJ. Dysregulation of the TOX-RUNX3 pathway in cutaneous T-cell lymphoma. Oncotarget. 2015;10(33):3104–3113.

28. Thonnart N, Caudron A, Legaz I, et al. KIR3DL2 is a coinhibitory receptor on Sézary syndrome malignant T cells that promotes resistance to activation-induced cell death. Blood. 2014;124(22):3330–3332.

29. Lay L, Stroup B, Payton JE. Validation and interpretation of IGH and TCR clonality testing by Ion Torrent S5 NGS for diagnosis and disease monitoring in B and T cell cancers. Pract Lab Med. 2020;22:e00191.

30. Borcherding N, Bormann NL, Kraus G. scRepertoire: An R-based toolkit for single-cell immune receptor analysis. F1000Res. 2020;9:47.

31. Bates SE, Eisch R, Ling A, et al. Romidepsin in peripheral and cutaneous T-cell lymphoma: mechanistic implications from clinical and correlative data. Br. J. Haematol. 2015;170(1):96–109.

32. Lopez AT, Bates S, Geskin L. Current Status of HDAC Inhibitors in Cutaneous T-cell Lymphoma. Am J Clin Dermatol. 2018;19(6):805–819.

33. Lu G, Jin S, Lin S, et al. Update on histone deacetylase inhibitors in peripheral T-cell lymphoma (PTCL). Clinical Epigenetics. 2023;15(1):124.

34. Wang P, Wang Z, Liu J. Role of HDACs in normal and malignant hematopoiesis. Mol Cancer. 2020;19(1):5.

35. Whittaker SJ, Demierre M-F, Kim EJ, et al. Final Results From a Multicenter, International, Pivotal Study of Romidepsin in Refractory Cutaneous T-Cell Lymphoma. Journal of Clinical Oncology. 2010;28(29):4485–4491.

36. Beygi S, Fernandez-Pol S, Duran G, et al. Pembrolizumab in mycosis fungoides with PD-L1 structural variants. Blood Adv. 2021;5(3):771–774.

37. Khodadoust MS, Rook AH, Porcu P, et al. Pembrolizumab in Relapsed and Refractory Mycosis Fungoides and Sézary Syndrome: A Multicenter Phase II Study. JCO. 2020;38(1):20–28.

38. Horwitz SM, Koch R, Porcu P, et al. Activity of the PI3K-δ,γ inhibitor duvelisib in a phase 1 trial and preclinical models of T-cell lymphoma. Blood. 2018;131(8):888–898.

39. Horwitz SM, Moskowitz AJ, Jacobsen ED, et al. The Combination of Duvelisib, a PI3K-δ,γ Inhibitor, and Romidepsin Is Highly Active in Relapsed/Refractory Peripheral T-Cell Lymphoma with Low Rates of Transaminitis: Results of Parallel Multicenter, Phase 1 Combination Studies with Expansion Cohorts. Blood. 2018;132(Suppl 1):683–683.

40. Beygi S, Duran GE, Fernandez-Pol S, et al. Resistance to mogamulizumab is associated with loss of CCR4 in cutaneous T-cell lymphoma. Blood. 2022;139(26):3732–3736.

41. Jameson KL, Mazur PK, Zehnder AM, et al. IQGAP1 scaffold-kinase interaction blockade selectively targets RAS-MAP kinase-driven tumors. Nat Med. 2013;19(5):626–630.

42. Choi S, Hedman AC, Sayedyahossein S, et al. Agonist-stimulated phosphatidylinositol-3,4,5-trisphosphate generation by scaffolded phosphoinositide kinases. Nat Cell Biol. 2016;18(12):1324–1335.

43. Thines L, Roushar FJ, Hedman AC, Sacks DB. The IQGAP scaffolds: Critical nodes bridging receptor activation to cellular signaling. Journal of Cell Biology. 2023;222(6):e202205062.

44. Hodge RG, Ridley AJ. Regulating Rho GTPases and their regulators. Nat Rev Mol Cell Biol. 2016;17(8):496–510.

45. Etienne-Manneville S, Hall A. Rho GTPases in cell biology. Nature. 2002;420(6916):629–635.

46. Kiel MJ, Sahasrabuddhe A a, Rolland DCM, et al. Genomic analyses reveal recurrent mutations in epigenetic modifiers and the JAK-STAT pathway in Sézary syndrome. Nature communications. 2015;6:8470.

47. Netchiporouk E, Litvinov IV, Moreau L, et al. Deregulation in STAT signaling is important for cutaneous T-cell lymphoma (CTCL) pathogenesis and cancer progression. Cell Cycle. 2014;13(21):3331–3335.

48. Béguelin W, Teater M, Meydan C, et al. Mutant EZH2 Induces a Pre-malignant Lymphoma Niche by Reprogramming the Immune Response. Cancer Cell. 2020;37(5):655–673.e11.

49. Béguelin W, Popovic R, Teater M, et al. EZH2 is required for germinal center formation and somatic EZH2 mutations promote lymphoid transformation. Cancer cell. 2013;23(5):677–92.

50. Bödör C, Grossmann V, Popov N, et al. EZH2 mutations are frequent and represent an early event in follicular lymphoma. Blood. 2013;122(18):3165–8.

51. Goldsmith SR, Fiala MA, O’Neal J, et al. EZH2 Overexpression in Multiple Myeloma: Prognostic Value, Correlation With Clinical Characteristics, and Possible Mechanisms. Clin Lymphoma Myeloma Leuk. 2019;19(11):744–750.

52. Li B, Chng W-J. EZH2 abnormalities in lymphoid malignancies: underlying mechanisms and therapeutic implications. Journal of Hematology & Oncology. 2019;12(1):118.

53. Morin RD, Johnson NA, Severson TM, et al. Somatic mutations altering EZH2 (Tyr641) in follicular and diffuse large B-cell lymphomas of germinal-center origin. Nature Genetics. 2010;42(2):181–185.

54. van Kemenade FJ. Coexpression of BMI-1 and EZH2 polycomb-group proteins is associated with cycling cells and degree of malignancy in B-cell non-Hodgkin lymphoma. Blood. 2001;97(12):3896–3901.

55. Velichutina I, Shaknovich R, Geng H, et al. EZH2-mediated epigenetic silencing in germinal center B cells contributes to proliferation and lymphomagenesis. Blood. 2010;116(24):5247–55.

56. Dasgupta M, Dermawan JKT, Willard B, Stark GR. STAT3-driven transcription depends upon the dimethylation of K49 by EZH2. Proc Natl Acad Sci U S A. 2015;112(13):3985–3990.

57. Kim E, Kim M, Woo D-H, et al. Phosphorylation of EZH2 activates STAT3 signaling via STAT3 methylation and promotes tumorigenicity of glioblastoma stem-like cells. Cancer Cell. 2013;23(6):839–852.

58. Viré E, Brenner C, Deplus R, et al. The Polycomb group protein EZH2 directly controls DNA methylation. Nature. 2006;439(7078):871–874.

59. Kim J, Lee Y, Lu X, et al. Polycomb- and Methylation-Independent Roles of EZH2 as a Transcription Activator. Cell Rep. 2018;25(10):2808–2820.e4.

60. Souroullas GP, Jeck WR, Parker JS, et al. An oncogenic Ezh2 mutation induces tumors through global redistribution of histone 3 lysine 27 trimethylation. Nature Medicine. 2016;22(6):632–640.

61. Yan J, Li B, Lin B, et al. EZH2 phosphorylation by JAK3 mediates a switch to noncanonical function in natural killer/T-cell lymphoma. Blood. 2016;128(7):948–958.

62. Zimmerman SM, Lin PN, Souroullas GP. Non-canonical functions of EZH2 in cancer. Front Oncol. 2023;13:1233953.

63. Zimmerman SM, Nixon SJ, Chen PY, et al. Ezh2Y641F mutations co-operate with Stat3 to regulate MHC class I antigen processing and alter the tumor immune response in melanoma. Oncogene. 2022;41(46):4983–4993.

64. Batlevi CL, Salles G, Park SI, et al. Tazemetostat in Combination with Lenalidomide and Rituximab in Patients with Relapsed/Refractory Follicular Lymphoma: Phase 1b Results of Symphony-1. Blood. 2022;140(Supplement 1):2296–2298.

65. Duan R, Du W, Guo W. EZH2: a novel target for cancer treatment. Journal of Hematology & Oncology. 2020;13(1):104.

66. Izutsu K, Ando K, Nishikori M, et al. Tazemetostat for relapsed/refractory B-cell non-Hodgkin lymphoma with EZH2 mutation in Japan: 3-year follow-up for a phase II study. Int J Hematol. 2024;

67. Izutsu K, Makita S, Nosaka K, et al. An open-label, single-arm phase 2 trial of valemetostat for relapsed or refractory adult T-cell leukemia/lymphoma. Blood. 2023;141(10):1159–1168.

68. McCabe MT, Ott HM, Ganji G, et al. EZH2 inhibition as a therapeutic strategy for lymphoma with EZH2-activating mutations. Nature. 2012;492(7427):108–112.

69. Morschhauser F, Tilly H, Chaidos A, et al. Tazemetostat for patients with relapsed or refractory follicular lymphoma: an open-label, single-arm, multicentre, phase 2 trial. Lancet Oncol. 2020;21(11):1433–1442.

70. Olsen EA, Kim YH, Kuzel TM, et al. Phase IIb multicenter trial of vorinostat in patients with persistent, progressive, or treatment refractory cutaneous T-cell lymphoma. J. Clin. Oncol. 2007;25(21):3109–3115.

71. Piekarz RL, Frye R, Turner M, et al. Phase II Multi-Institutional Trial of the Histone Deacetylase Inhibitor Romidepsin As Monotherapy for Patients With Cutaneous T-Cell Lymphoma. Journal of Clinical Oncology. 2009;27(32):5410–5417.

72. Schade AE, Kuzmickas R, Rodriguez CL, et al. Combating castration-resistant prostate cancer by co-targeting the epigenetic regulators EZH2 and HDAC. PLOS Biology. 2023;21(4):e3002038.

