## Supplemental Methods for "Single Cell Resolution Tracking of Cutaneous T-Cell Lymphoma Reveals Clonal Evolution in Disease Progression"

### Peripheral blood mononuclear cell isolation, purification and flow cytometry/sorting

PBMCs were subjected to enrichment and/or depletion using antibody cocktails (Miltenyi Biotec) to enable purification of the desired cells. For depletion of monocytes, peripheral blood samples were incubated with RosetteSep Human Monocyte (CD36) Depletion Cocktail for 15 minutes at room temperature, layered onto a Histopaque-1077 gradient, and centrifuged at 400g for 30 minutes with no brake. The interphase was collected and washed with 10 mL of sort buffer (PBS, 1% FBS, 2 mM EDTA), followed by red blood cell lysis (155 mM $NH_4Cl$, 10 mM $KHCO_3$, 0.1 mM EDTA) for 10 minutes. Cells were washed and resuspended in sort buffer. For purification of T cell populations, the EasySep Human CD4 Positive Selection Kit was used according to the manufacturer's instructions, or cells were stained with CD4 (Miltenyi Biotec Cat 130-092-373, RRID:AB_871684), CD7 (Miltenyi Biotec Cat 130-105-842, RRID:AB_2659107), and/or CD26 (Miltenyi Biotec Cat 130-093-441, RRID:AB_1103210) fluorophore conjugated antibodies and subjected to FACS on a Sony iCyt Synergy SY3200. For isolation of NK cells, monocyte- and T-cell depleted PBMCs were stained with the above antibodies and anti-CD56 antibody (Miltenyi Biotec AF12-7H3 Cat 130-113-309) and subjected to FACS on a Sony iCyt Synergy SY3200. Alternatively, PBMCs were isolated from peripheral blood using a Histopacque-1077 gradient followed by red blood cell lysis as described(Andrews et al., 2019) and then cryopreserved for future purification. If cryopreserved, PBMCs were suspended in 90% FBS with 10% DMSO and placed overnight in a -80°C freezer using a Mr. Frosty container (ThermoFisher Scientific cat. 5100-0001) to achieve an optimal rate of cooling, then transferred to liquid nitrogen storage.

### Flow sorting for live CD4+ cells from peripheral blood mononuclear cells for scRNA-seq

Cryopreserved cells ($2\text{-}10 \times 10^6$) were thawed as follows: cryovials of cells were warmed in a 37°C water bath until the cell began to thaw. After ~1 minute, 1 mL of ambient temperature fetal bovine serum (FBS) was added to the cryovial and allowed to mix before removal to a fresh tube. This process was repeated until the cells were fully thawed. Thawed cells were pelleted by centrifugation at 400g for 5 minutes and resuspended in sort buffer (phosphate-buffered saline (PBS) + 2% FBS + 2 mM EDTA) at a concentration of $1 \times 10^7$/mL. A subset of cells were then stained for CD4 (Miltenyi Biotec Cat 130–092-373, RRID:AB_871684), 7-AAD (Miltenyi Biotec Cat 130–111-568), and CD3 (Miltenyi Biotec Cat 130-113-128, RRID:AB_2725956), fluorophore conjugated antibodies according to manufacturer instructions and subjected to flow cytometry. The remainder of the cells were 7-AAD (Miltenyi Biotec Cat 130–111-568) and viable cells were sorted by FACS on a Sony iCyt Synergy SY3200 to collect live cells. These cells were immediately processed for scRNAseq (see below).

### Exome + whole genome (eWGS) sequencing

Exome + whole-genome sequencing (eWGS) was performed on DNA isolated from purified malignant T cells or matched non-malignant cells from peripheral blood, or from frozen or FFPE biopsies (skin or lymph node). In total, there were 26 samples from ten MF/SS patients. Genomic DNA (450-600ng for WGS, 50-250 ng for exome-seq) was fragmented on the Covaris LE220 instrument targeting 375bp (WGS) or 250bp (exome) inserts. Automated libraries were constructed with the KAPA Hyper PCR-free library prep kit (KAPA Biosystems/Roche) on the SciClone NGS platform (Perkin Elmer). For WGS, the fragmented genomic DNA was size selected on the SciClone instrument with AMPure XP beads to tighten the distribution of fragmented DNA to ensure the average insert of the libraries were 350-375bp. We followed the manufacturer's protocol as provided by Perkin Elmer, with the following exception: Post ligation, the libraries were purified twice with a 0.7x AMPure bead/sample ratio to eliminate any residual adaptors present. An aliquot of the final libraries was diluted 1:5 and quantitated on the Caliper GX instrument (Perkin Elmer). For exome-seq, 1 low input manual amplified library (input 23ng) was made using the Swift Accel-NGS 2S DNA Library Kit – 96rxn. Ten libraries were pooled at an equimolar ratio yielding ~4.5µg per library pool prior to the hybrid capture. The library pool was hybridized with the xGen Lockdown Exome Panel v1.0 reagent (IDT Technologies). The concentration of each library was accurately determined through qPCR utilizing the KAPA library Quantification Kit according to the manufacturer's protocol (KAPA Biosystems/Roche) to produce cluster counts appropriate for the Illumina NovaSeq6000 instrument. Normalized libraries were sequenced on a NovaSeq6000 S4 Flow Cell using the XP workflow and a 151x10x10x151 sequencing recipe according to manufacturer protocol. Target coverage was 30Gb per sample for WGS and 12Gb per sample for exome-seq.

#### eWGS data processing and analysis

Exome and whole genome sequencing Fastq files were aligned with bwa to hg38, duplicate reads marked with PicardTools, and unmapped and duplicate reads filtered from the analysis with samtools (Li et al., 2009; "Picard Tools - By Broad Institute," n.d.) . For copy number analysis, GATK Mutect2 v4.2.0.0 (Van der Auwera and O'Connor, 2020) was used to call germline variants to determine B-allele frequency (BAF); these variant calls were used to determine loss of heterozygosity or allelic imbalance due to genomic copy number changes. CNVkit(Talevich et al., 2016) was used to call copy number alterations (CNA) in 100kb windows using germline variants to calculate BAF, and segmentation was performed using default circular binary segmentation. CNVkit genemetrics was used to find genes associated with CNA. Any CNA falling within blacklist regions on hg38 were removed from the analysis. CNA with log2 fold change <abs(0.2) were excluded. CNA in second or third samples were compared to patient-matched first samples. If a CNA was not detected in a patient's first sample, but was detected in subsequent samples, it was considered to be Acquired with Progression or Resistance to therapy (APR).

For somatic single nucleotide variant (sSNV) analysis, GATK Mutect2 v4.2.0.0 (Van der Auwera and O'Connor, 2020) was used to call somatic variants in exome sequencing data by comparing each tumor sample to the matched normal with subsequent filtering with FilterMutectCalls. Somatic SNVs that passed the Mutect filter were annotated by VEP(McLaren et al., 2016) and predictions for variant effect were made by SIFT, PolyPhen, CONDEL, and LoFtool(Adzhubei et al., 2010; Clifford et al., 2004; Fadista et al., 2017; Ng and Henikoff, 2003). Non-synonymous moderate to high impact somatic sSNVs with ≥20 variant-containing reads and ≥100 total reads covering the sSNV site were retained for downstream analyses. C>T sSNVs detected only in FFPE samples were determined to be likely artifacts and removed(Do and Dobrovic, 2015). If a sSNV passing these filters was not detected in a patient's first sample, but was detected in subsequent samples, it was considered to be APR.

#### Single cell RNA and VDJ sequencing (scRNA-seq)

For each of 16 PBMC samples, 10,000 to 20,000 viable cells were submitted for processing using the 10x Genomics Chromium Controller and the Chromium Single Cell 5′ Library & Gel Bead Kit v2 (PN-1000006) following the manufacturer's protocols. cDNA was prepared after the GEM generation and barcoding, followed by the GEM-RT reaction and bead cleanup steps. Purified cDNA was amplified for 10-14 cycles before being cleaned up using SPRIselect beads. Samples were then run on a Bioanalyzer to determine the cDNA concentration. TCR target enrichments were performed on the full-length cDNA. GEX and Enriched TCR libraries were prepared as recommended by the 10x Genomics Chromium Single Cell V(D)J Reagent Kits (v1.0 Chemistry) user guide with appropriate modifications to the PCR cycles based on the calculated cDNA concentration. For sample preparation on the 10x Genomics platform, the Chromium Single Cell 5 'Library and Gel Bead Kit v2 (PN-1000006), Chromium Single Cell A Chip Kit (PN-1000152), Chromium Single Cell V(D)J Enrichment Kit, Human, Tcell (96rxns)(PN-1000005), and Chromium Single Index Kit T (PN-1000213) were used. The concentration of each library was accurately determined through qPCR utilizing the KAPA library Quantification Kit according to the manufacturer's protocol (KAPA Biosystems/Roche) to produce cluster counts appropriate for the Illumina NovaSeq6000 instrument. Normalized libraries were sequenced on a NovaSeq6000 S4 Flow Cell using the XP workflow and a 151x10x10x151 sequencing recipe according to manufacturer protocol. A median sequencing depth of 50,000 reads/cell was targeted for each sample.

**STAT3 functional assays**

*Luciferase assay*: These were performed as we have before(Milner et al., 2015) (. Briefly, STAT3-null A4 cells (derived from DLD cells, kindly provided by Dr G. Stark, Cleveland Clinic, Cleveland, OH)(Yang et al., 2010) were transfected with plasmids encoding STAT3_Wild Type, STAT3_LOF (K558E), STAT3_GOF (N647I), or STAT3_p.D661Y expression vectors and STAT3-driven firefly luciferase reporter and control renilla reporter plasmids per the manufacturer's instructions (Cignal STAT3; Qiagen). After 24 hours, IL-6 (10ng/mL) was added to cells, control cells received PBS. After another 24 hours, luciferase activity was assayed as a measure of STAT3-driven transcription, followed by measurement of firefly and renilla luciferase (Dual-Luciferase Reporter Assay; Promega). Results were calculated as the ratio of reporter (firefly) to control (renilla) luciferase. Cytokine stimulation was conducted in serum-containing media.

*GFP-reporter assay*: The STAT3-GFP reporter cell line was generated by stable transduction of a lentiviral STAT3 reporter construct (Addgene plasmid #110495) into a STAT3-deficient mammalian A4 cell line (DLD cells)(Yang et al., 2010), via infection at a multiplicity of infection (MOI) 20. The reporter construct consists of four copies of a transcriptional response element (TRE), facilitating phosphorylated STAT3 dimer binding, which drives the expression of green fluorescent protein (GFP). The construct was packaged into lentivirus by the Hope Center Viral Vectors Core at Washington University School of Medicine. For reporter assays, $6x10^5$ cells/mL of the STAT3-GFP reporter cell line were seeded in 6-well polystyrene tissue culture plates. When 70% confluent, they were transiently transfected with WT or *STAT3* mutants (STAT3_LOF (K558E), STAT3_GOF (N647I), or STAT3_p.D661Y) plasmids (as above) using Lipofectamine 3000 transfection reagent mixture (Thermo Fisher Scientific, USA) according to the manufacturer's instructions. After 24 hours, transfected STAT3-GFP reporter A4 cells were stimulated with IL-6 (10ng/mL, 24 hours) and GFP expression was measured by flow cytometry.

**STAT3 CUT&RUN-sequencing**

STAT3-null A4 cells(Yang et al., 2010) transfected with plasmids encoding STAT3_Wild Type or STAT3_p.D661Y expression vectors were subjected to CUT&RUN essentially as in(Michael P Meers et al., 2019; Skene and Henikoff, 2017) using modifications to the Cell Signaling Technology (CST) CUT&RUN assay kit (catalog #86652). Briefly, one hundred thousand cells per antibody were incubated with activated concanavalin magnetic beads plus antibody (anti-STAT3, CST 91395, or isotype and species matched IgG, Millipore Sigma 12-371) the solution incubated with rotation overnight at 4 degrees C. The next day, pAG-MNase enzyme in digitonin buffer was added and the solution incubated with rotation for 1 hour at 4 degrees C. DNA was then digested, isolated, and purified per the manufacturer's instructions. Isolated DNA was subjected to library preparation and clean-up using standard procedures(Michael P Meers et al., 2019; Skene and Henikoff, 2017) using unique i5/i7 primers and sequenced using an Illumina MiSeq 150 cycle kit paired end 2x75bp yielding an average of 22 million reads per run. Alignment and peak calling were performed as we have previously(Cubitt et al., 2024), based on(Michael P. Meers et al., 2019). Briefly, fastq files were aligned to GRCh38 using bowtie2 and samtools, reads trimmed using trimmomatic, bedgraphs generated using bedtools, and peaks called using SEACR(Langmead and Salzberg, 2012; Li et al., 2009; Michael P. Meers et al., 2019; Quinlan and Hall, 2010). ChIPseeker(Wang et al., 2022; Yu et al., 2015) was used to annotate peaks to functional genomic regions; GSEA was used to identify enriched pathways(Reimand et al., 2019).

**scRNA-seq data processing and analysis**

Alignment and gene counting were performed using the Cell Ranger pipeline (10x Genomics, v3.0, Pleasanton, CA). Genes found in fewer than 15 cells in each sample were removed. For each patient, gene counts and cells were pooled into a single Seurat (v5.1.0)(Butler et al., 2018; Hao et al., 2024; Stuart et al., 2019) object, and cells containing fewer than 200 or more than 3000-3750 expressed genes, more than 8-10% mitochondrial reads, fewer than 300 or more than 10000UMIs, or classification as a doublet by the R package scDblFinder(Germain et al., 2022) with parameters dims = 30, clust.method = "fast_greedy" were removed. In total, 118,608 cells were sequenced, with 92,496 passing initial QC filters after removing dead cells, empty droplets, and suspected multiplets. Normalization and regression of technical variation due to mitochondrial read percentage and read depth was performed with the SCTransform function(Choudhary and Satija, 2022; Hafemeister and Satija, 2019) with variables.features.n = 4000. Integration to account for experimental variability due to differences in preparation and batch effects was performed using the Seurat wrapper around the fastMNN function from the batchelor R package (v1.4.0)(Haghverdi et al., 2018) with n.features = 3000. Gene expression was normalized and the top 1500 variable using the "VST" method were calculated. Immune receptor genes were manually removed from the variable gene list before scaling the data and calculating the top 30 principal components. Data was integrated using the harmony (v1.0.0)(Korsunsky et al., 2019) R package using both patient and batch information to correct for batch effect with up to 10 iterations. The UMAP and neighbors were calculating with Seurat, using 20 dimensions of the harmony calculations. Unbiased differential expression analysis was performed with Seurat FindAllMarkers to identify genes with expression profiles that defined each of the 24 expression clusters.

**scRNA-seq annotation of cell type**

Cell annotation was performed using the singleR (v1.4.1)(Aran et al., 2019, n.d.) R package with the HPCA(Mabbott et al., 2013) data set as references and the fine label discriminators. Individual sequencing runs were subsetted to run through the singleR algorithm in order to reduce memory demands. The output of all the singleR analyses were collated and appended to the meta data of the Seurat object. Cell type designations with less than 50 cells in the entire cohort were reduced to "other". Automated annotations were checked manually using canonical marker genes.

**scRNA-seq TCR repertoire analysis**

The filtered contig annotation T cell receptor (TCR) data for available sequencing runs were loaded into the R global environment. Individual contigs were combined using the combineTCR() function of scRepertoire (v1.3.2)(Borcherding et al., 2020) R Package. Clonotypes were assigned to barcodes and multiple duplicate chains for individual cells were filtered to select for the top expressing contig by read count. The clonotype data was then added to the Seurat Object using combineExpression with proportion across individual patients (including all the timepoints of sequencing) being used to calculate frequency.

#### scRNA-seq copy number estimation

Single-cell copy number estimates were calculated using the CopyKat (v1.0.5)(Gao et al., 2021) R package using default settings. The normal cells, as defined as singleR annotation of monocytes, T or NK Cells and non-expanded clonotypes (proportion of repertoire < 0.01 or < 0.1 depending on patient), were used. In the absence of reference normal cells or malignant cells, copy number was not estimated. Estimated aneuploid classification from CopyKat was combined with the single-cell meta data and copy number estimates to generate UMAPs across multiple timepoints.

#### Detection of sSNVs in scRNAseq data

For each patient, the filtered list of sSNVs detected by analysis of exome sequencing as described above was compared to scRNAseq datasets for that patient using cb_sniffer(Lambo et al., 2023; Petti et al., 2019; sridnona, 2024). For each sSNV, cell barcodes with sequencing reads containing the variant were retrieved. The following filters were then applied: sSNVs with sequencing reads from ≥20 unique cell barcodes per sample in ≥1 sample, cell barcode variant allele frequency ≥ 0.20 (per sample = number of unique cell barcodes containing sSNV / number of total unique cell barcodes).

#### Integrating and prioritizing CNA and sSNV

For defining and prioritizing recurrently mutated genes and mutations that were acquired with progression or therapy resistance in this cohort for highlighting in Figures 3, 6, and S6, the following filters were applied:

- ≥1 CNA and ≥1 sSNV across all samples, total of ≥2 patient IDs.
- ≥1 APR CNA or sSNV, total of ≥2 patient IDs.
- >100 basemean expression level in MF/SS samples (bulk RNAseq)
- Detected in ≥1 patient ID in scRNAseq data

#### Single cell visualization

All single cell visualizations were performed with Seurat v5.1.0 and ggplot2 R package v3.5.1.
